# *Plasmodium falciparum* exploits NUAK1 to establish infection in human erythrocytes

**DOI:** 10.1101/2025.10.30.685469

**Authors:** Daniel J. Navarrete, Chi Yong Kim, Kyle McLelland, Mario Gonzalez Ramirez, Barbara Baro, Nichole D. Salinas, Niraj H. Tolia, Christian Doerig, Shao-En Ong, Martin G. Golkowski, Elizabeth S. Egan

## Abstract

The malaria parasite *Plasmodium falciparum* continues to demonstrate growing drug resistance, raising the need for innovative treatments. Host-directed therapeutics are emerging as a promising approach for many infectious diseases, but knowledge of critical host factors for malaria is limited. *P. falciparum* is an obligate intracellular parasite of human erythrocytes, suggesting it has evolved to exploit specific host pathways to establish infection. Here, we report that the AMPK-related kinase NUAK1 is a critical host factor for *P. falciparum* in erythrocytes and has potential as a therapeutic target. We show that NUAK1 is present in human erythrocytes and undergoes increased phosphorylation in *P. falciparum*-infected cells. Two highly selective NUAK1 inhibitors, HTH-01-015 and WZ4003, inhibited *P. falciparum* growth throughout its asexual life cycle, including during erythrocyte invasion. Chemoproteomic profiling confirmed the inhibitors’ selectivity for human NUAK1. We further show that treatment with the inhibitors reduces phosphorylation of the well-characterized NUAK1 substrate MYPT1 in erythroid cells. Moreover, we find that genetic overexpression of NUAK1 in erythroid cells partially rescues both the signaling and invasion phenotypes elicited by the small molecule inhibitors. These results establish a critical role for the NUAK1 signaling pathway in *P. falciparum*-infected erythrocytes and highlight its potential as a vulnerable target for host-directed malaria control.

## INTRODUCTION

Despite effective antiparasitic drugs, the malaria parasite *Plasmodium falciparum* continues to cause nearly 600,000 deaths and hundreds of millions of symptomatic cases every year ^1^. One long-standing challenge to malaria control is the parasite’s proclivity to develop drug resistance ^2^, a pattern that has been repeatedly observed throughout history ^3–5^. Most recently, *P. falciparum* parasites with mutations that confer resistance to the highly potent first-line antimalarial agent, dihydroartemisinin, have begun to arise in Southeast Asia and sub-Saharan Africa, highlighting an urgent need for novel treatment approaches ^5–9^.

Host-directed therapies offer an advantage in treating infectious diseases by depriving the pathogen of the most direct way to develop resistance, namely selection of mutations in the drug target, and by providing opportunities to target novel pathways ^10,11^. Such host-directed therapies have proven effective for a variety of infectious diseases, such as tuberculosis, HIV, and COVID-19 ^12–15^; however, the potential of host-targeted therapies for malaria has so far received only limited attention ^16,17^. Erythrocyte (red blood cell, RBC) invasion is a key step in the asexual life cycle of *P. falciparum* and involves a series of intricate protein-protein interactions between the parasite and host cell. For example, the erythrocyte binding-like (EBL) family of *P. falciparum* proteins are critical for proper attachment of the invasive merozoite onto the RBC in preparation for the subsequent steps of invasion ^18–20^. One EBL protein, EBA-175, binds to RBC plasma membrane protein glycophorin A (GYPA) ^21,22^; in addition to being important for attachment, this interaction has been shown to modulate the phosphorylation of erythrocyte cytoskeletal proteins and increase the deformability of the host cell, presumably promoting invasion ^23–26^. Recent efforts identified erythrocyte CD44 as an additional binding partner for EBA-175, and showed that it likely acts as a co-receptor during invasion by facilitating signaling to the host cell cytoskeleton ^27,28^.

Here, we used phospho-antibody microarrays to investigate host signaling pathways that may be stimulated during *P. falciparum* invasion of human RBCs and identified the AMPK-related kinase NUAK1 as a candidate host factor activated during invasion. Using genetic and small molecule-based approaches, we demonstrate that the NUAK1 signaling pathway is active in human RBCs and provide experimental evidence that direct targeting of this host pathway inhibits *P. falciparum* infection. We propose that NUAK1 signaling plays a critical role in erythrocyte cytoskeletal homeostasis, which is key to *P. falciparum* invasion. This work highlights the potential for selective targeting of a human kinase in the context of host-directed antimalarial intervention.

## RESULTS

### Human NUAK1 is phosphorylated during *P. falciparum* infection

To investigate the CD44-dependent signaling induced by EBA-175, we generated isogenic wild-type (WT) or CD44-null cultured red blood cells (cRBCs) from primary human hematopoietic stem/progenitor cells (HSPCs) and used phospho-antibody microarrays produced by Kinexus (Fig. 1a). This method has previously been employed to investigate host protein phosphorylation in *P. falciparum*-infected erythrocytes ^29^. Analysis of the microarray data identified NUAK1 as an intriguing candidate, as increased signal for anti-NUAK1 pThr211 was detected for WT cRBCs stimulated with EBA-175 or rupturing schizonts, but not for CD44-null cRBCs stimulated in the same manner (Extended Data Fig. 1). This observation was validated through dedicated immunoblotting, which showed increased phosphorylation of NUAK1 at Thr211 in lysates from WT cRBCs stimulated with EBA-175, but not in CD44-null cRBCs (Fig. 1b).

**Figure 1.**
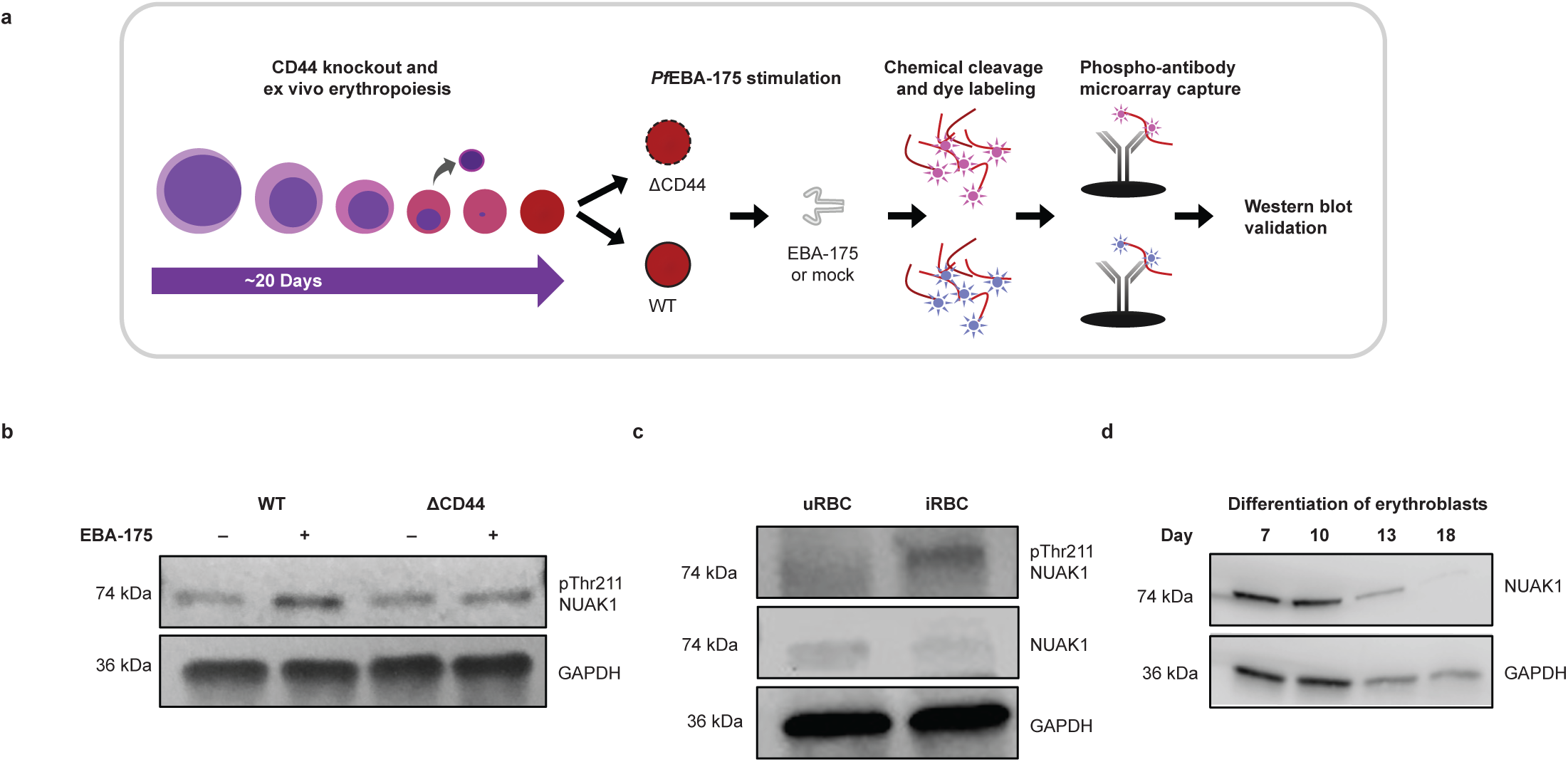
Stimulation with EBA-175 or *P. falciparum* infection increases NUAK1 phosphorylation in RBCs. **a**, Schematic representation of ex-vivo erythropoiesis protocol to generate isogenic WT and CD44-null cRBC from primary human CD34+ HSPCs, followed by Kinexus phosphoantibody microarrays. **b**, Western blot of NUAK1 Thr211 phosphorylation in whole cell lysates from WT and CD44-null mature cRBCs (day 21) +/-EBA-175 stimulation. **c**, Western blot of NUAK1 Thr211 in RBC supernatant fractions from uninfected (uRBC) and *P. falciparum*-infected RBCs (iRBC). **d**, Western blot of NUAK1 expression in primary human CD34+ HSPC-derived erythroblasts at indicated dates of ex-vivo differentiation.

NUAK1, or ARK5, is a serine/threonine kinase and a member of the AMPK-related kinase family. Overexpression of NUAK1 is associated with a variety of human cancers, where its activation can promote cell survival under conditions of oxidative stress as well as cell migration and the epithelial mesenchymal transition, driving metastasis ^30–36^. NUAK1 is canonically activated by the tumor suppressor LKB1 via phosphorylation of Thr211 ^37^, and its major substrate is the myosin phosphatase complex, MYPT1. To our knowledge, NUAK1 has not previously been characterized in erythrocytes.

To determine if *P. falciparum* infection can activate NUAK1 phosphorylation in mature RBCs from peripheral blood, we compared the amount of NUAK1 pThr211 in lysates from *P. falciparum*-infected RBCs (iRBCs) versus uninfected RBCs (uRBCs) by immunoblotting. We observed increased phosphorylation of NUAK1 (pThr211) in iRBCs compared to uRBCs, similar to the results for EBA-175-stimulated cRBCs (Fig. 1c). These findings are also consistent with phospho-antibody microarrays performed by Adderley *et al.* (2020), which showed increased NUAK1 phosphorylation in lysates from *P. falciparum*-infected RBCs at the ring, trophozoite, and schizont stages, as compared to uninfected RBCs ^29^. To determine the dynamics of NUAK1 expression within the erythroid lineage, we performed western blots on CD34^+^ HSPCs induced to undergo ex-vivo erythropoiesis and observed a decrease in protein abundance during terminal erythroid differentiation (Fig. 1d). Together, these results demonstrate that NUAK1 is expressed in erythroid cells, establish the presence at low levels of NUAK protein in mature erythrocytes, and indicate that the NUAK1 signaling pathway may play a role in *P. falciparum* infection.

### A potential role for NUAK1 during erythroid proliferation

To investigate the potential role of human NUAK1 in erythrocytes, we first attempted a knock-out approach in primary human CD34^+^ HSPCs using nucleofection of ribonucleoprotein complexes containing sgRNAs targeting NUAK1 and recombinant Cas9 protein. However, after inducing the transfected cells to differentiate down the erythroid lineage, we failed to recover NUAK1-null cRBCs, suggesting NUAK1 may have an essential function during erythroid differentiation. As an alternative approach, we employed two compounds that have been shown to exhibit high selectivity for NUAK: WZ4003, which inhibits both NUAK1 and NUAK2, and HTH-01-015, which is specific for NUAK1 ^38^. We found that both HTH-01-015 and WZ4003 inhibited proliferation of the BEL-A immortalized erythroid cell line, which resembles proerythroblasts ^39^ (Fig. 2a). These results suggest that the small molecules HTH-01-015 and WZ4003 have a target in the erythroid lineage– potentially NUAK1, and that its inhibition is detrimental to the survival of erythroid lineage cells.

**Figure 2.**
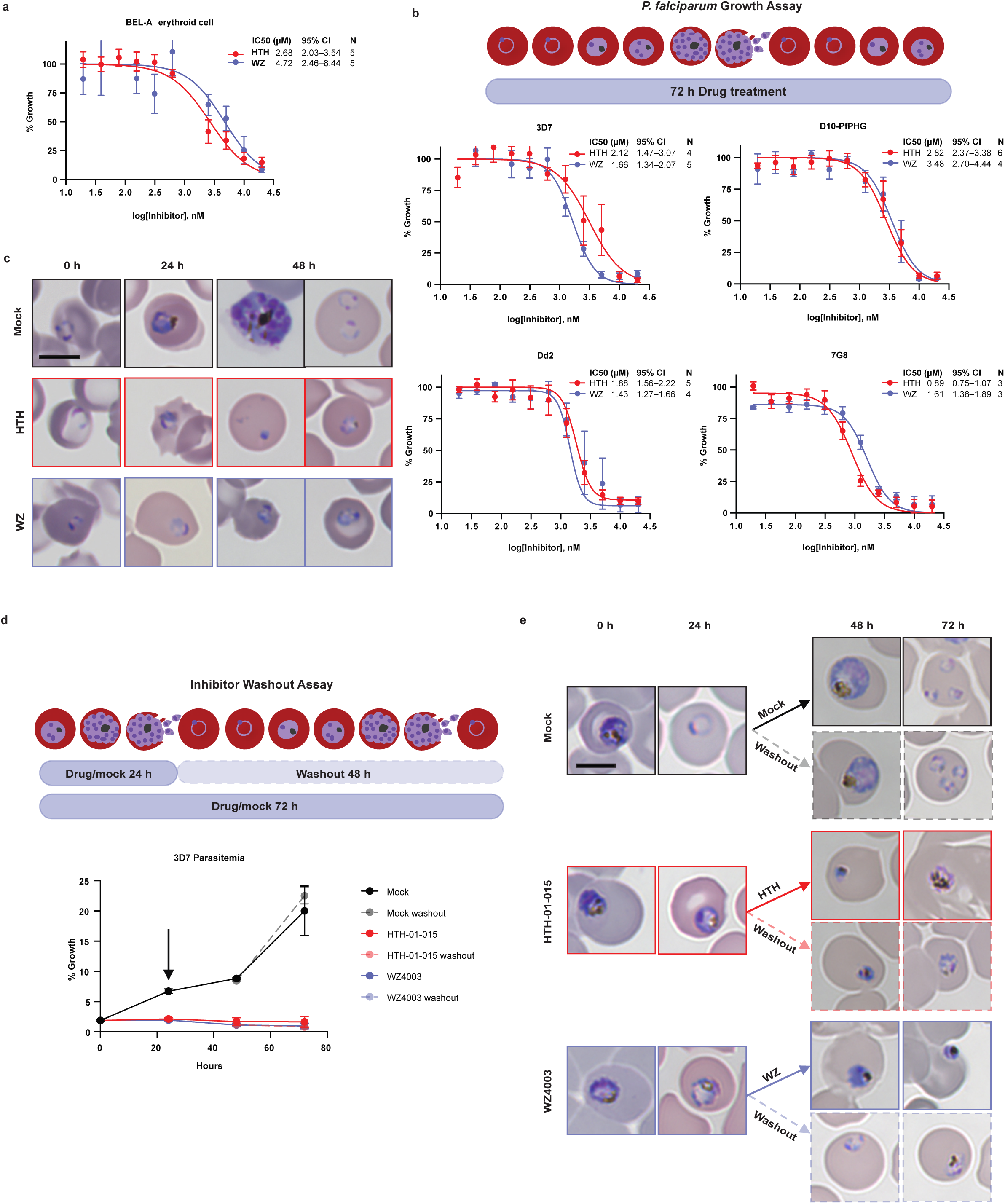
HTH-01-015 and WZ4003 inhibit erythroid cell proliferation and *P. falciparum* growth in vitro. **a**, Dose-response curves for BEL-A cells cultured in the presence of NUAK1 inhibitors HTH-01-015 or WZ4003 for 72 hours. Cell concentration was measured by flow cytometry and plotted relative to vehicle control. Mean ± SEM, N = 5 biological replicates. IC50 values and 95% confidence intervals (CI) are shown. **b**, Dose-response curves for four lab-adapted *P. falciparum* strains cultured for 72 hours with HTH-01-015 or WZ4003. Parasitemia was measured by flow cytometry and plotted relative to vehicle control. Mean ± SEM, N = 3–6 biological replicates, as indicated. IC50 values and 95% CI are shown. **c**, Giemsa-stained cytospins of *P. falciparum* strain 3D7 time course treated with 5 µM of each indicated inhibitor. Scale bar = 5 µm. Mock indicates vehicle control. **d**, Growth curve of inhibitor washout assay. *P. falciparum* 3D7 was incubated with 5 µM of inhibitor or vehicle control beginning at the trophozoite stage (0 hours), either washed to remove drug or maintained in drug at 24 hours (arrow) and cultured until 72 hours. Parasitemia was measured every 24 hours by flow cytometry. Mean ± SEM, N = 2. **e**, Representative Giemsa-stained cytospins from the washout experiment. Scale bar = 5 µm.

### NUAK1 inhibitors have antimalarial activity

To investigate the effects of the NUAK1 inhibitors on *P. falciparum*, we incubated ring-stage parasites with increasing concentrations of each compound and measured the parasitemia after 72 hours. We found that each of the four lab-adapted *P. falciparum* strains tested, 3D7, D10-PfPHG, 7G8, and Dd2, was inhibited by both HTH-01-015 and WZ4003 in a dose-dependent manner, with IC_50_ values in the low micromolar range (Fig. 2b). Examination of Giemsa-stained cytospins prepared from a time course of drug exposure revealed that *P. falciparum* parasites exposed to inhibitor at the ring stage failed to develop past the early trophozoite stage (Fig. 2c). To determine if the parasites could recover after drug treatment, we treated trophozoite-stage parasites with 5 µM of NUAK1 inhibitors or mock (vehicle control) for 24 hours before washing off the drug. Assessment of parasitemia by flow cytometry revealed that mock-treated parasites grew 10-fold in 72 hours while inhibitor-treated parasites did not increase, even after washing out the drug (Fig. 2d). Giemsa-stained cytospins revealed pyknotic trophozoites in the inhibitor-treated conditions by 48 hours, regardless of washout (Fig. 2e). These data indicate that the compounds have cidal, rather than static, antimalarial activity during the asexual intraerythrocytic cycle.

### NUAK1 inhibitors block *P. falciparum* invasion

We next aimed to examine whether the NUAK1 inhibitors specifically block invasion of erythrocytes by *P. falciparum*. Prior to this stage of the parasite life cycle, mature schizonts rupture, releasing invasive merozoites, which then can attach to and invade nearby RBCs via a multi-step process involving erythrocyte deformation, discharge of the rhoptry organelles, and formation of a moving junction, followed by internalization. To test if the NUAK1 inhibitors can specifically inhibit invasion, we quantified merozoite egress and invasion in the presence of HTH-01-015, WZ4003, mock (vehicle control), or ML10, a well-characterized inhibitor of *P. falciparum* PfPKG that inhibits egress ^40^. Parasitemia was measured by flow cytometry at 0 and 3 hours to capture the changes in the population as schizonts ruptured and new rings were established (Fig. 3a).

**Figure 3.**
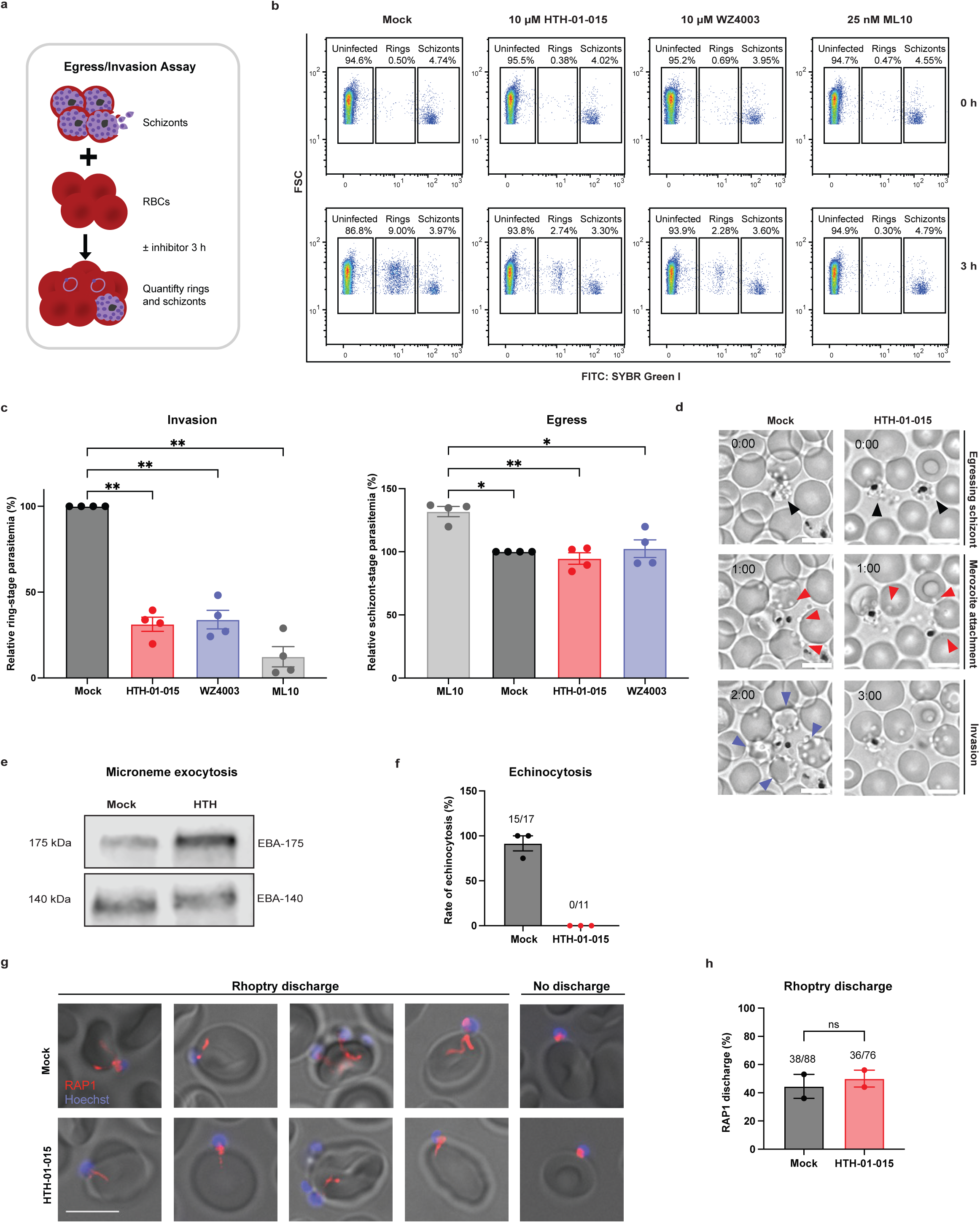
NUAK1 inhibitors block *P. falciparum invasion* of RBCs. **a,** Schematic representation of the egress/invasion assay. **b**, Flow cytometry plots from one representative egress/invasion assay showing ring- and schizont-stage parasitemia at 0 and 3 hours as quantified using SYBR Green I. **c**, Results from egress/invasion assays at T = 3 hr, showing ring-stage (Invasion) and schizont-stage (Egress) parasitemia in the presence of each drug, normalized to vehicle control (Mock). Mean ± SEM, N = 4. Each dot represents a biological replicate. Statistical analysis: Ordinary one-way ANOVA; *p < 0.05, **p < 0.01. **d**, Live microscopy images of a time course of *P. falciparum* egress and invasion in presence of 10 µM HTH-01-015 or mock (vehicle control). Black arrows indicate rupturing schizonts, red arrows indicate merozoites attached to RBCs, and blue arrows indicate invaded RBCs. Scale bar = 8 µm. **e**, Western blot of EBA-175 and EBA-140 expression in culture supernatants from invading *P. falciparum* merozoites. **f**, Frequency of RBC echinocytosis induced by attached merozoites in the presence of HTH-01-015 or Mock, as measured by live microscopy. All samples included Cyt-D to prevent internalization. Each point represents an independent experiment, with the denominator indicating the total number of merozoite-RBC pairs scored (attached > 30 s) and the numerator indicating those in which echinocytosis was observed. **g**, Representative immunofluorescence assay images for PfRAP1 localization (red) in RBCs with attached merozoites (Hoechst, blue). Scale bar = 5 µm. **h**, Frequency of rhoptry discharge from attached merozoites as measured by RAP1 localization within the RBC displayed as the mean ± SEM, N = 2 independent experiments, with the denominator indicating the total number of merozoite-RBC pairs scored and the numerator indicating those in which RAP1 discharge was observed in the RBC. Statistical analysis: Two-tailed, two-sample Student’s t-test; “ns”, not significant.

In the mock condition, we observed an expected drop in the schizont-stage parasitemia between 0 and 3 hours, with a concomitant rise in the ring-stage parasitemia to up to 9%, reflecting schizont rupture and robust merozoite invasion in the absence of inhibitors (Fig. 3b,c). In contrast, *P. falciparum* invasion was significantly inhibited in the presence of either HTH-01-015 or WZ4003, as indicated by the low percentage of ring stage parasites at 3 hours relative to mock (< 3% versus ∼9%) (Fig. 3b,c). Notably, the schizonts decreased to a similar degree in the mock and the NUAK1 inhibitor conditions, demonstrating that the observed invasion phenotype was not due to a block in schizont rupture. In comparison, as expected schizonts failed to rupture and egress in the presence of the egress inhibitor control, ML10. These results were further confirmed by live-cell imaging, where we observed schizont egress and merozoite attachment in the presence of the NUAK1 inhibitors, but failed to detect invasion events (Fig. 3d, Extended Data Videos 1 and 2). Together, these results show that the NUAK1 inhibitors HTH-01-015 and WZ4003 specifically inhibit *P. falciparum* invasion of human RBCs.

Why is invasion blocked in the presence of the NUAK inhibitors? After schizont egress, *P. falciparum* merozoites release EBA and Rh family proteins from the microneme organelles, which aid in host cell attachment, apical reorientation, and erythrocyte deformation^20,41^. This process of microneme exocytosis is stimulated by an increase in calcium within the parasite, and results in re-localization of the invasion ligands onto the apical surface of the merozoite. We tested whether HTH-01-015 inhibits microneme exocytosis using an established assay that measures proteins released into the media in a culture of merozoites attempting to invade RBCs. Both EBA-175 and EBA-140 were readily detected in the spent media, regardless of the presence of the NUAK1 inhibitor (Fig. 3e). These findings demonstrate that microneme exocytosis is not affected by the NUAK1 inhibitors and are consistent with the idea that these compounds are likely inhibiting invasion by acting on a host target.

The next step of invasion involves release of the parasite’s rhoptry organelle contents into the host cell, which requires an interaction between the merozoite invasion ligand PfRH5 and Basigin on the RBC^42,43^. Rhoptry release has been proposed to induce a transient phenotype of RBC echinocytosis observable by live-cell imaging, as echinocytosis is blocked by antibodies that inhibit the interaction between PfRH5 and Basigin ^19,44^. To test whether the NUAK inhibitor HTH-01-015 impacts echinocytosis, we used an established live imaging assay that quantifies echinocytosis in the presence of the actin polymerization inhibitor cytochalasin-D, which prevents merozoite internalization without affecting merozoite attachment or rhoptry discharge. We observed a striking difference in the rate of echinocytosis for the two conditions, with ∼90% of RBCs with an attached merozoite observed to undergo echinocytosis in the control condition, versus 0% in the presence of HTH-01-015 (Fig. 3f, Extended Data Videos 3 and 4).

To directly test whether this echinocytosis phenotype was due to failure of rhoptry discharge, we used immunofluorescence assays to visualize the rhoptry bulb protein RAP1 during the process of merozoite invasion in the presence or absence of HTH-01-015. If rhoptry discharge were impaired, RAP1 would be expected to remain within the attached merozoite, rather than be injected into the RBC. We observed a similar rate of rhoptry discharge in both the control and HTH-01-015-treated conditions, as indicated by ∼50% of merozoite-RBC attachment pairs displaying “whorls” of RAP1 localized within the RBC (Fig. 3g,h). These results indicate that HTH-01-015 does not inhibit rhoptry discharge, despite inhibiting echinocytosis, consistent with the implication of a host cell factor. Moreover, these findings suggest that the lack of merozoite-induced echinocytosis and internalization in the presence of this drug is attributable to an effect on the host cell, rather than the parasite.

### HTH-01-015 and WZ4003 are selective to human NUAK1

In vitro kinase assays have demonstrated high selectivity of HTH-01-015 and WZ4003 for NUAK1, relative to 140 other human kinases, including several AMPK family members ^38^. As these inhibitors have not been previously studied in *P. falciparum* nor in erythroid cells, we turned to an unbiased approach for target identification. We employed Kinobead competition assays, a chemical proteomic approach in which sepharose beads coated with broadly promiscuous kinase inhibitors are used to enrich diverse kinases from a protein lysate ^45^. Incubating native cell lysates with ATP-competitive kinase inhibitors leads to the competition for target binding between the Kinobeads and the inhibitor, as revealed by quantitative mass spectrometry analyses of enriched kinases, comparing inhibitors versus vehicle controls ^46^.

Initial attempts to perform Kinobead assays with human erythrocytes were unsuccessful, as the abundant hemoglobin in these cells bound to the beads nonspecifically, leading to saturation. We next turned to a mixed mammalian cell lysate, reasoning that use of cells expressing many kinases would strengthen confidence in the results if a specific target were identified (Fig. 4a). Of the 263 human kinases that were enriched by the Kinobeads, only two kinases were significantly competed for binding by HTH-01-015 and WZ4003: NUAK1 and cyclin G-associated kinase (GAK) (Fig. 4b,c, Extended Data Table 1). HTH-01-015 and WZ4003 competed for NUAK1 with log_2_ differences of 2.1 (p< 0.01) and 5.7 (p< 0.001), respectively, and GAK, with log_2_ differences of 1.2 (p< 0.001) and 2.0 (p< 0.001), respectively. These results confirm the high selectivity of both HTH-01-015 and WZ4003 for NUAK1, even within a diverse cellular lysate. While GAK was identified as an additional potential target of these compounds, this kinase has previously been reported to bind to a variety of kinase inhibitors, suggesting the interaction may be low affinity off-target ^47,48^. To test this idea, we performed in vitro growth assays to evaluate a highly selective GAK inhibitor, 12G ^49^, for its effect on *P. falciparum*. Notably, the IC_50_ value for 12G was approximately ten times higher than those of HTH-01-015 or WZ4003 (14.7 µM versus 1.8 µM and 1.9 µM, respectively), suggesting that the antimalarial phenotypes of HTH-01-015 and WZ4003 are unlikely to be due to GAK inhibition (Extended Data Fig. 2).

**Figure 4.**
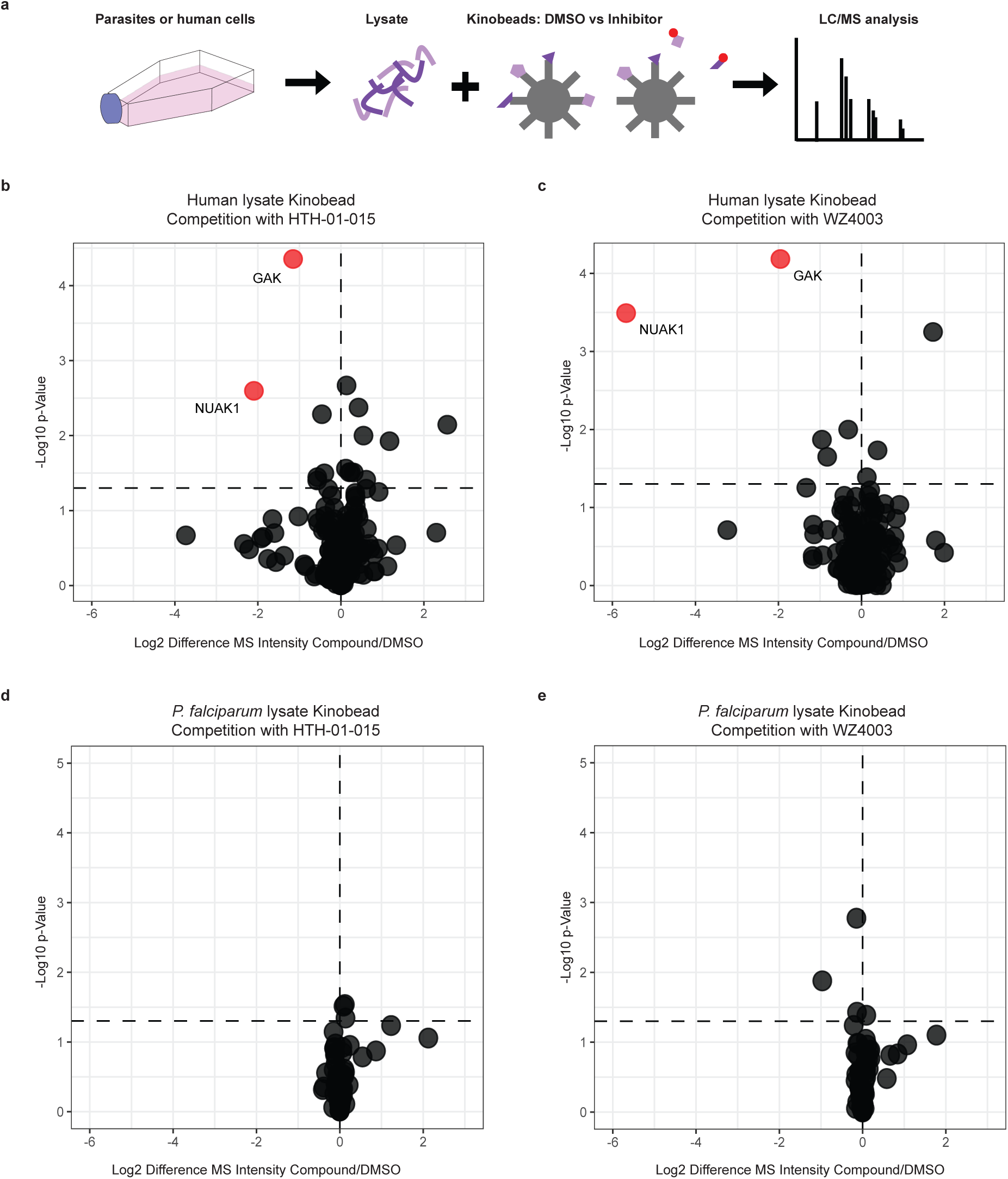
Kinobead competition assays identify human NUAK1 as top target of HTH-01-015 and WZ4003. **a**, Schematic representation of Kinobead kinase inhibitor competition assay. **b-c**, Volcano plots comparing Kinobead enrichment from mixed human cell lysate (hepatocyte and Jurkat cells) in presence of each NUAK1 inhibitor (50 μM) versus DMSO control. 263 unique human kinases were identified. N = 3. Significantly competed kinases (Log2 fold change < -1 and p-value < 0.05) are shown in red and non-competed kinases in black. **d-e**, Volcano plots comparing Kinobead enrichment from *P. falciparum* lysate (60 unique parasite kinases) in presence of each NUAK1 inhibitor (50 μM) versus DMSO control. Statistical analysis: Significantly competed kinases were determined using a two-tailed two-sample Student’s t-test with Benjamini-Hochberg correction for multiple hypothesis testing.

Although bioinformatic searches using publicly available databases failed to identify a *P. falciparum* orthologue of NUAK1, we next wanted to directly test if the antimalarial activity of HTH-01-015 or WZ4003 may be due to an effect on a parasite kinase. We again turned to Kinobead competition assays as an unbiased approach. As an apicomplexan parasite, *P. falciparum* has a reduced kinome relative to mammalian cells yet is known to express ∼90 protein kinases ^50–54^. We have shown previously that kinobeads can be used to profile kinase inhibitor selectivity in Apicomplexan parasites ^54^. Notably, of the 60 parasite protein kinases that were enriched from a *P. falciparum* lysate, none of them were significantly competed by either of the NUAK1 inhibitors (Fig. 4d, e), while 10 µM of the non-selective control competitor staurosporine significantly competed 18 parasite kinases (Extended Data Table 1).

As another approach to investigate potential parasite targets, we utilized a split-luciferase-based kinase screening assay developed by Luceome that included a custom panel of 11 *Plasmodium* kinases alongside human NUAK1 and NUAK2 ^55,56^. HTH-01-015 and WZ4003 both displayed high binding activity to NUAK1, as measured by ∼85%-95% reduction in luciferase activity (Extended Data Fig. 3a, b). Of the *P. falciparum* kinases in the panel, PfNEK3 and PfPKB showed moderate interactions with one or both inhibitors (neither of these kinases was detected in the Kinobeads assays), but both of these kinases have been shown previously to be nonessential: PfNEK3 was found to not be required for asexual-stage development of *P. falciparum* ^57^ and both PfNEK3 and PfPKB were found to be dispensable in a *piggyBac* transposon mutagenesis screen, with high mutability index scores (0.6, 0.98, respectively) ^58^. Together, the results from the Kinobead and split-luciferase assays demonstrate that the most likely target of HTH-01-015 and

WZ4003 in *P. falciparum*-infected erythrocytes is NUAK1, and that the antimalarial activity of these compounds is due to inhibition of this host kinase.

### NUAK1 signaling pathway is functional in erythroid cells

As our findings pointed to human NUAK1 as the most likely target of HTH-01-015 and WZ4003 in *P. falciparum*-infected erythrocytes, we next sought to functionally characterize the NUAK1 pathway in erythroid cells. To do this, we employed the BEL-A erythroid cell line, which can be induced to differentiate down the erythroid lineage, offering the opportunity to genetically characterize host factors in erythroid cells and their role in *P. falciparum* invasion ^39,59^.

In other cells, activated NUAK1 is known to phosphorylate MYPT1 at the Ser445 residue, and treatment with NUAK1 inhibitors leads to reduced phosphorylation ^35,38,60^. To functionally interrogate the NUAK1-MYPT1 signaling pathway in the BEL-A cells, we incubated them with HTH-01-015, WZ4003, or vehicle control, and performed immunoblotting to detect MYPT1 as well as the phosphorylated form of the protein. We found that MYPT1 phosphorylation at Ser445 was indeed detectable in the BEL-A lysate, suggesting that the NUAK1 signaling pathway is present and active in these cells (Fig. 5a). Furthermore, we found that treatment with the NUAK1 inhibitors ablated the MYPT1 pSer445 signal, demonstrating that MYPT1 Ser445 phosphorylation in BEL-A cells is sensitive to NUAK1 inhibition (Fig. 5a,b).

**Figure 5.**
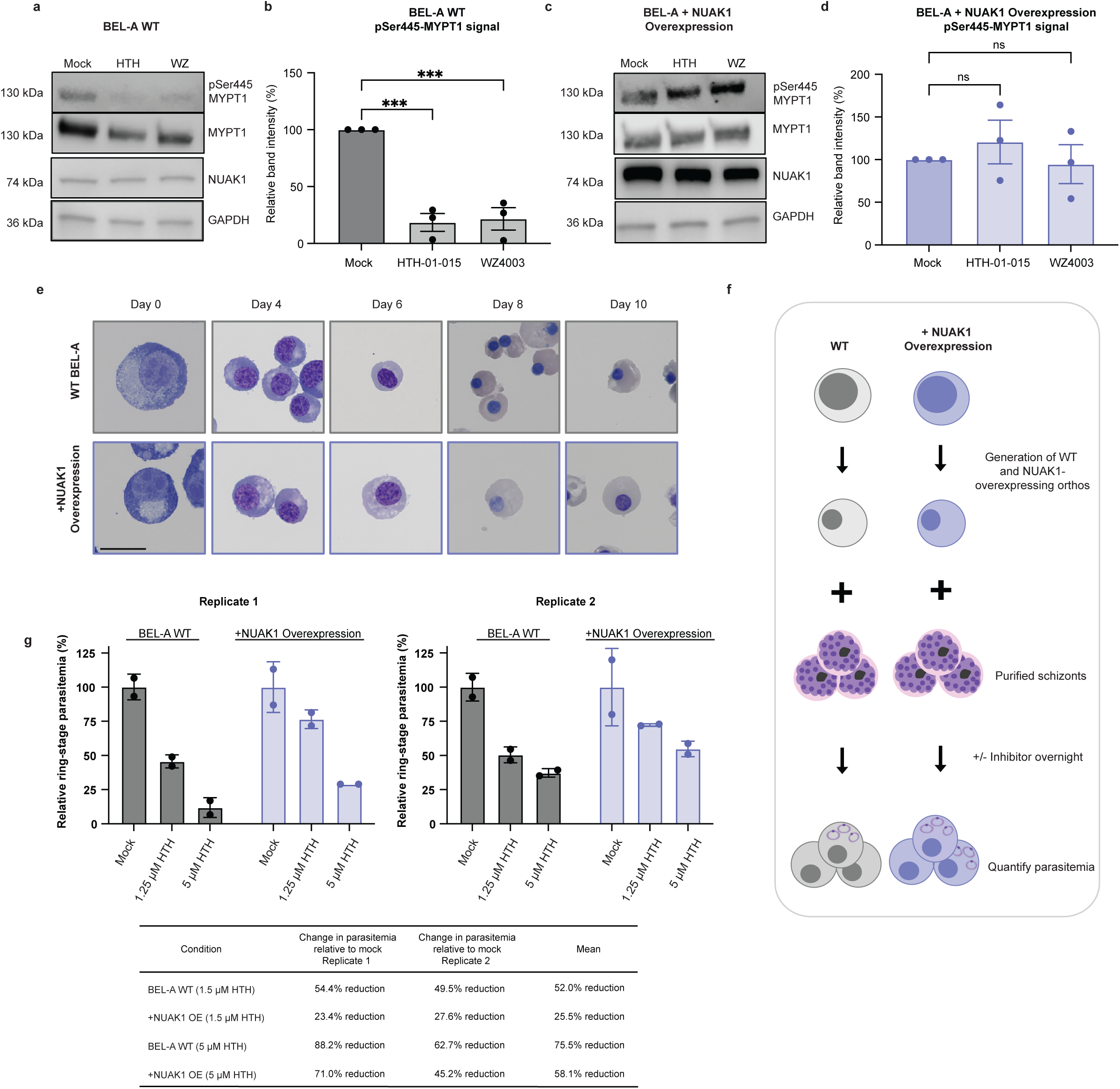
NUAK1 overexpression counteracts the inhibitory effects of HTH-01-015 on MYPT1 phosphorylation and *P. falciparum* invasion. **a, c,** Representative western blots of MYPT pSer445, total MYPT1, NUAK1, and GAPDH in WT BEL-A cells (**a**) or BEL-A overexpressing NUAK1 (**c**) after overnight treatment with 10 µM HTH-01-015, 10 µM WZ4003, or mock (vehicle control). **b, d,** Quantification of western blot results from three independent experiments. Band intensity of MYPT1 pSer445 was normalized relative to total MYPT1 signal, then plotted relative to the mock condition for each experiment. Mean ± SEM, N = 3. Statistical analysis: One-way ANOVA; *** p < 0.001. **e**, Giemsa-stained cytospin slides of WT and NUAK1-overexpressing BEL-A cells during erythroid differentiation. Scale bar = 20 µm. **f**, Schematic depicting *P. falciparum* invasion assays using orthochromatic erythroblasts (orthos) derived from WT or NUAK1-overexpressing BEL-A cells ± HTH-01-015. **g**, Results of *P. falciparum* strain 3D7 invasion assays in WT and NUAK1-overexpressing BEL-A orthos ± HTH-01-015. Mean ± SD. Each graph shows data from one independent experiment with points representing technical replicates.

To further validate the functionality of NUAK1 in the erythroid lineage and demonstrate the selectivity of the NUAK1 inhibitors, we generated a BEL-A cell line overexpressing NUAK1 using lentiviral transduction. After confirming overexpression of NUAK1, we incubated the cells with HTH-01-015, WZ4003, or vehicle control, and performed immunoblotting to evaluate MYPT1 phosphorylation in the presence or absence of the inhibitors. In contrast to the WT BEL-A cells, the MYPT1 pSer445 signal was preserved in BEL-A cells overexpressing NUAK1 even when they were treated with NUAK1 inhibitors, indicating that NUAK1 overexpression rescued the inhibitory phenotype (Fig. 5c,d). These experiments confirm a functional NUAK1 signaling pathway in erythroid cells that is targeted by the selective inhibitors.

### NUAK1 overexpression rescues *P. falciparum* invasion in erythroid cells

Having demonstrated that overexpression of NUAK1 in BEL-A cells reduced their sensitivity to the NUAK1 inhibitors, we next investigated the potential for NUAK1-overexpression to rescue the inhibitory effect of HTH-01-015 on *P. falciparum* invasion. To this end, we induced WT BEL-A or BEL-A overexpressing NUAK1 to differentiate down the erythroid lineage to orthochromatic erythroblasts (orthos) (Fig. 5e). Using *P. falciparum* strain 3D7, we performed invasion assays on the differentiated BEL-A orthos from the two different genetic backgrounds in the presence of either HTH-01-015 or mock (vehicle control) (Fig. 5f). As expected, we found that invasion into the WT BEL-A orthos was significantly inhibited by HTH-01-015, with a reduction in parasitemia of 52% and 76% with 1.25 µM and 5µM HTH-01-015, respectively, compared to mock vehicle control (Fig. 5g,h). In comparison, the invasion inhibitory effect of HTH-01-015 in the BEL-A orthos overexpressing NUAK1 was much milder, with a reduction in parasitemia of only 26% and 58% with 1.25 µM and 5 µM HTH-01-015, respectively, compared to mock (Fig. 5g,h). Thus, overexpressing NUAK1 substantially rescues the *P. falciparum* invasion phenotype of HTH-01-015, providing strong genetic evidence that *P. falciparum* exploits human NUAK1 during erythrocyte invasion. Moreover, parasite invasion was notably higher in cells overexpressing NUAK1 (Extended Data Fig. 4), further highlighting the role of NUAK1 as a host factor for invasion.

## DISCUSSION

Host-directed therapies hold great potential to optimize the treatment of infectious diseases by opening new pathways to target while minimizing the chance of developing resistance. Indeed, emerging reports highlight the successful inhibition of host factors for diverse viral and bacterial pathogens ^10,14,15,61,62^. Although *P. falciparum* is an obligate intracellular parasite of red cells that must be critically reliant on host factors for survival, the identity, function, and vulnerability of such factors are only beginning to be explored ^13,63,64^. Recent studies have investigated the clinical utility of SYK kinase inhibitors, which are proposed to block *P. falciparum* egress by inhibiting band 3 phosphorylation ^65–67^. Adderley *et al.* (2020) assessed signaling in *P. falciparum*-infected RBCs more broadly using phospho-antibody microarrays, leading to the discovery that a MAPK kinase pathway involving the receptor tyrosine kinase C-MET is activated in infected cells, and the identification of several inhibitors of this pathway with antimalarial activity ^29^. Yet despite these insights, to our knowledge no study has experimentally demonstrated inhibitor-selectivity for a specific human kinase in *P. falciparum*-infected RBCs, leaving open the possibility that the antimalarial activities of these inhibitors may be due to off-target effects on either host or parasite proteins.

In this study, we uncovered a role for the AMPK-related kinase NUAK1 in the invasion of human erythrocytes by *P. falciparum*. NUAK1 is upregulated in many cancers, where it is canonically activated by the tumor suppressor LKB1 and acts to promote cell migration and protect tumor cells from oxidative damage through its interaction with the myosin phosphatase complex ^34^. While NUAK1 has not previously been described in the red cell proteome, our research revealed its expression throughout erythropoiesis and demonstrates an active NUAK1-MYPT1 signaling pathway in erythroid cells. These insights open new questions about the roles of NUAK1 throughout hematopoiesis, including during normal development and malignant transformation, which are yet to be explored.

Our findings indicate that two selective small-molecule inhibitors of NUAK1, HTH-01-015 and WZ4003, have antimalarial activity throughout the parasite’s asexual lifecycle, including specifically during RBC invasion. *P. falciparum* invasion has been shown to involve changes in the phosphorylation of host cell cytoskeletal proteins as well as altered cell membrane deformability, but the host signaling pathways and molecular mechanisms driving successful invasion are poorly understood. We found that HTH-01-015 did not impede microneme exocytosis nor merozoite attachment to the RBC surface, but did inhibit RBC echinocytosis, a transient invasion-related phenotype that has been attributed to release of the rhoptry contents. However, direct visualization of the rhoptry bulb protein RAP1 in invading merozoites revealed that it was injected into the RBC with similar efficiency in the presence or absence of the inhibitor. Critically, these results demonstrate that echinocytosis can be uncoupled from rhoptry discharge, and, together with our chemoproteomic and genetic studies implicating NUAK1 as the most likely target of HTH-01-015, strongly suggest that both echinocytosis and internalization are dependent on the host NUAK1 signaling pathway. In other cells, activation of the NUAK1-MYPT1 pathway regulates cytoskeletal motor proteins that promote cell detachment ^35,38,68^. We found that treatment with the NUAK1 inhibitors led to reduced MYPT1 phosphorylation in the BEL-A erythroid cell line and that NUAK1 overexpression rescued MYPT1 signaling in the presence of the inhibitors. Further characterization of these inhibitors through global phosphoproteomics of *P. falciparum*-infected RBCs may illuminate novel pathways and interactions between the host cell and parasite during infection.

In conclusion, our findings reveal a role for the NUAK1 signaling pathway during *P. falciparum* infection of red blood cells. By demonstrating NUAK1’s importance in erythrocyte invasion and establishing host kinase selectivity of the small molecule inhibitor, we open new avenues for the generation of effective host-directed antimalarial therapeutics that may reduce the risk of developing drug resistance.

## MATERIALS AND METHODS

### *P. falciparum* culture and drug assays

1. *P. falciparum* strains were routinely grown in de-identified human erythrocytes (Stanford Blood Center) in complete RPMI (cRPMI), consisting of RPMI 1640 with 25 mM N-2-hydroxyethylpiperazine-N′-2-ethanesulfonic acid, 50 mg/L hypoxanthine, 2.42 mM sodium bicarbonate, and 0.5% Albumax (Invitrogen) at 37°C in 5% CO_2_ and 1% O_2_. Strains 3D7, Dd2, and 7G8 are lab-adapted strains obtained from the Walter and Eliza Hall Institute (Melbourne, Australia) and MR4 (BEI Resources). Strain D10-PfPHG was provided by Dave Richard ^69^.
2. *P. falciparum* drug assays were initiated with synchronized ring-stage *P. falciparum* starting at 0.5-1% parasitemia in 1% hematocrit in 120 µL volume in 96-well plates and incubated with serial dilutions of the inhibitors or DMSO as vehicle control for 72 hours in triplicate. Inhibitors used: HTH-01-015 (Sigma, #SML1446), WZ4003 (Sigma, #SML1445), and GAK inhibitor 12G (Sigma, #5.38770). Upon harvest, cells were stained in SYBR Green I (1:2,000, Invitrogen #S7563) in PBS/0.3% BSA for 20 minutes and events were collected by flow cytometry using a MACSQuant flow cytometer (Miltenyi) to determine the parasitemia. The D10-PfPHG-infected samples were washed in PBS/0.3% BSA and run on the flow cytometer without SYBR Green staining as they express GFP. For each assay, we performed a minimum of three independent biological replicates. For the inhibitor washout assays, synchronous trophozoite-stage cultures were incubated with 5 µM HTH-01-015, 5 µM WZ4003, or mock vehicle control (0.05% DMSO) for 24 hours at ∼2% parasitemia and 1% hematocrit in 120 µL volume in a 96-well plate in triplicate. Cultures were then washed and either resuspended in drug-free media or in their respective drug treatment. Parasitemia was measured by flow cytometry every 24 hours until the 72-hour endpoint. For the washout assay, we performed two independent biological replicates. Samples were additionally analyzed by microscopy of cytospins stained with Giemsa and May-Grünwald.

### BEL-A cell culture, differentiation, and drug assays

BEL-A cells were obtained from NHS Blood and Transplant (NHSBT) and routinely plated at a concentration of 1×10^5^ cells/mL in proliferation media consisting of StemSpan (Stemcell Technologies) with 100 ng/mL SCF (R&D Systems), 3 U/mL erythropoietin (Epo; Amgen), 10^-6^ M dexamethasone (Sigma), and 1 µg/mL doxycycline (Sigma) at 37°C in 5% CO_2_ and split every two days ^39^. To induce differentiation, cells were transitioned to erythroid differentiation medium composed of Iscove Basal Medium (IMDM) (Sigma, #FG0465) supplemented with 4 mM L-Glutamine (Sigma), 330 µg/ml holo-transferrin (BBI Solutions), 10 µg/ml of recombinant human insulin (Sigma), 2 IU/ml heparin (Fisher), 0.5% v/v Pen-Strep (Fisher), 10 ng/ml SCF, 3 IU/ml Epo, 1 µg/mL doxycycline, and 5% human plasma (Octapharma) at a concentration of 5×10^5^ cells/mL. On day 2 of differentiation, doxycycline was removed from the media and cells were kept at 1 × 10^6^ cells/mL. On day 4 SCF was removed, cells were maintained at 1 × 10^6^ cell/mL until day 8 for harvest/assay setup. Cell stage was monitored by cytospin preparations stained with May-Grünwald and Giemsa.

BEL-A drug assays were initiated at 1×10^5^ cells/mL in proliferation media in 96-well plates at 120 µL/well in serial dilutions of HTH-01-015, WZ4003, or DMSO as vehicle control. After 72 hours, cell concentrations were measured using a MACSQuant flow cytometer, and growth was plotted relative to the DMSO condition.

### Egress-Invasion assays

Late-stage *P. falciparum* schizonts were isolated by magnetic separation using a MACS magnet (Miltenyi Biotec) and incubated in 25 nM of ML-10 (LifeArc) in cRPMI for up to 3 hours. After three washes to remove ML-10, the synchronized schizonts were incubated with erythrocytes at a starting parasitemia of 4% and 1% hematocrit in the presence of 10 µM HTH-01-015, 10 µM WZ4003, 25 nM ML-10, or 0.1% DMSO (mock vehicle control) in a 96-well plate (100 µL/well) in triplicate. Samples were harvested at 0 and 3 hours post-treatment. Parasitemia was quantified by flow cytometry after staining cells for 20 minutes with SYBR Green I (1:2,000 dilution in PBS/0.3% BSA). The ring-stage and schizont-stage parasitemias at 3 hours were calculated relative to the parasitemias of the mock vehicle control. The ML-10-treated sample was used as negative control for egress for comparison across conditions.

### Live cell imaging

For live imaging, late-stage magnet-purified schizonts were treated with 25 nM of ML-10 for 3.5 hours, washed and incubated with erythrocytes at ∼8% parasitemia at 0.25% hematocrit in 500 µL cRPMI in a 35 mm FluoroDish (WPI), treated with 10 µM HTH-01-015 or mock treatment (0.1% DMSO), and imaged using a Keyence BZ-X700 microscope equipped for live imaging at 37°C with 60X objective. For echinocytosis assays, cultures were treated with 1 µM cytochalasin-D to prevent parasite internalization. Rate of echinocytosis was calculated as the number of echinocytosis events observed per attached merozoite-RBC pair surface for ≥ 30 seconds for each of three independent experiments.

### Antibodies used

Commercial antibodies used for western blotting include rabbit anti-NUAK1 pThr211 (Kinexus, #PK737), rabbit anti-NUAK1 (Proteintech, #22723-1-AP), mouse anti-MYPT1 (Santa Cruz, #514261), and rabbit anti-GAPDH (Abcam, #ab9485). Anti-EBA-175 is rabbit serum (R1454) raised against a hexa his-tagged 3D7 EBA-175 fusion protein and anti-EBA-140 is rabbit serum (R228) raised against a GST-tagged EBA-140 R3-5 fusion protein; both were obtained from the Cowman lab. Anti-MYPT1 phospho-Ser 445 (S508B) is a sheep antibody which was obtained from MRC PPU Reagents, along with the non-phosphopeptide ^35^. Secondary antibodies used for western blotting included HRP-conjugated goat anti-mouse IgG (Cell Signaling, #7076), goat anti-rabbit IgG (Cell Signaling, #7074), and mouse anti-sheep IgG (Rockland, #18-8815-31). Antibodies used for immunofluorescence assays include mouse monoclonal anti-PfRAP1 209.3, a gift from Anthony Holder ^70^, and goat anti-mouse Alexafluor 647 secondary antibody (ThermoFisher, #A21235).

### Immunoblotting

All samples were resolved on SDS-PAGE gels (Bio-Rad), transferred to methanol-activated PVDF membranes using a Turbo Transfer system (Bio-Rad), and blocked in TBST (0.1% Tween 20) containing either 5% BSA or 5% non-fat dry milk (Bio-Rad, #1706404) for 1 hour at room temperature before primary antibody incubations. Blots were washed 3 times in TBST for 5 minutes in between antibody incubations. Specific lysis and antibody conditions for each experiment are detailed below. For all blots, chemiluminescent signal was developed using SuperSignal West Pico PLUS substrate (ThermoFisher, #34580).

For western blot validation of Kinexus phosphoantibody microarrays, 3.3 × 10^7^ day 21 CD44-null or WT cRBCs were incubated with 2 µM recombinant EBA-175 or mock (PBS) at 25% hematocrit for 30 minutes at 4_°_C. Whole cell lysates were prepared by resuspending the cells in 250 µL of Kinexus lysis buffer supplemented with protease inhibitors (Thermo, #A32955) and 1 µM DTT and incubating on ice for 10 minutes. After centrifugation, lysate supernatants were diluted 3:1 (v/v) with 4X Laemmli sample buffer (Bio-Rad, # 1610747), heated at 95_°_C for 5 minutes, and 7 µL of each diluted sample were loaded on an SDS-PAGE gel and prepared for western blotting. Following transfer, the membrane was blocked in TBST/5% BSA then cut at 50 kDa and the top piece was incubated in anti-NUAK1 pThr211 (1:1,000 in TBST/5% BSA) and the bottom piece was incubated in anti-GAPDH (1:8,000 in TBST/5% milk) antibodies at 4_°_C overnight. After washing, blots were incubated in anti-rabbit HRP secondary antibody (1:3,000 in TBST/5%BSA) at room temperature for one hour.

For western blotting of RBC lysates from infected vs. uninfected cells, lysates were prepared from 8 × 10^7^ magnet-purified trophozoites or 8 × 10^7^ uRBCs from the same blood donor. After washing, cells were resuspended in 300 µL 0.15% Saponin (Millipore) in PBS (lysate concentration of 2.7 × 10^8^ cells/mL) with 1X Halt protease and phosphatase inhibitor cocktail (ThermoScientific, #78440), on ice for 5 minutes. The samples were pelleted at 21,000 x g at 4_°_C and the supernatant (RBC fraction) was used for western blots while the pellet (parasite material) was discarded. Lysate supernatants were diluted 3:1 (v/v) in 4X Laemmli sample buffer containing 4% β-mercaptoethanol, heated at 95_°_C for 5 min, and 17 µL of each diluted sample was loaded on an SDS-PAGE gel in duplicate for western blotting. Following transfer, the PVDF membrane was blocked in TBST/5% BSA, then cut horizontally at the 50 kDa marker. The top half was then cut vertically to separate the duplicate sets of lanes. The two resulting top pieces were incubated separately, one with anti-NUAK1 pThr211 antibody (1:1,000 in TBST/5% BSA) and the other with anti-NUAK1 antibody (1:500 in TBST/5% BSA). The bottom piece was incubated with anti-GAPDH antibody (1:5,000 in TBST/5% BSA) at 4 °C overnight. Membranes were then incubated with anti-rabbit HRP secondary antibody (1:1,000 in TBST/5% BSA) at room temperature for 1 hour.

For western blots of primary human CD34^+^ HSPC-derived erythroblasts, cells at each day of differentiation were pelleted and resuspended in RIPA lysis buffer (ThermoScientific, #89900) at a concentration of 1 × 10^7^ cells/mL on ice for 10 minutes. Lysates were centrifuged at 21,000 x g at 4_°_C for 10 minutes and the supernatant was diluted 3:1 (v/v) in 4X Laemmli sample buffer containing 10% β-mercaptoethanol and heated at 95_°_C for 10 minutes. 15 µL of each diluted sample was loaded on a gel and prepared for western blotting. After transfer, the membrane was blocked in TBST/5% milk then cut at 50 kDa. The top piece was incubated in anti-NUAK1 (1:500 in TBST/5% milk) antibody and the bottom piece was incubated in anti-GAPDH (1:5,000 in TBST/5% milk) antibody at 4_°_C overnight, followed by incubation in anti-rabbit HRP secondary antibody (1:1,000 in TBST/5% milk) at room temperature for one hour.

For western blots of *P. falciparum* micronemal proteins, donor RBCs were treated with 3.334 mg/mL Chymotrypsin, 3.334 mg/mL Trypsin, and 0.5 u/mL Neuraminidase to remove surface receptors. Late-stage schizonts were magnet-purified and added to enzyme-treated RBCs at 8% parasitemia and 1% hematocrit in the presence of 10 µM HTH-01-015 or mock vehicle control (0.1% DMSO) in cRPMI for 3 hours at 37_°_C in 1% O_2_ and 5% CO_2_. 1 mL of supernatant was pelleted, and the supernatant was diluted 3:1 (v/v) in 4X Laemmli sample buffer containing 10% β-mercaptoethanol and heated at 95_°_C for 5 minutes. 17 µL of each diluted sample was loaded on to a gel and prepared for western blotting. After transfer, the membrane was blocked in TBST/5% milk, then cut vertically to separate the duplicate sets of lanes, then each piece was incubated with either anti-EBA-175 (1:500 TBST/5% milk) or anti-EBA140 (1:500 TBST/5% milk) rabbit serum at 4_°_C overnight, followed by incubation in anti-rabbit HRP secondary antibody (1:1,000 TBST/5% milk) at room temperature for one hour.

For BEL-A phospho-western blots, cells were incubated overnight with 10 µM HTH-01-015, 10 µM WZ4003, or mock vehicle control (0.1% DMSO) in proliferation media at 5 × 10^5^ cell/mL in a 6-well plate (2 mL/well). The following day, the cells were incubated with 100 nM of PP1/2 inhibitor, Calyculin A (ThermoFisher, #PHZ1044) for 30 min at 37_°_C before collection to preserve MYPT1 phosphorylation ^68,71^. Cells were pelleted and washed in PBS before being lysed in RIPA buffer with 1X Halt protease and phosphatase inhibitor cocktail and 100 nM Calyculin A at a concentration of 5 × 10^6^ cells/mL for 10 minutes on ice and centrifuged at 21,000 x g for 10 minutes. Lysate supernatant was diluted 3:1 (v/v) in 4X Laemmli sample buffer containing 10% β-mercaptoethanol, heated at 95_°_C for 5 minutes, and 15 µL of each diluted sample was prepared for western blotting as described above.

The sheep anti-MYPT1 pSer445 antiserum was pre-cleared prior to use by diluting the antiserum to 1 µg/mL in TBST/5% milk with the corresponding non-phospho-peptide (1:1,000 dilution) and incubating for 30 minutes at room temperature with rotation. To remove non-specific binding proteins, this mixture was then incubated with a blank, methanol-activated PVDF membrane (Bio-Rad, #1620174) for 1 hour. The resulting pre-cleared antibody solution was used for immunoblotting. After transfer, the membrane was cut at the 50 kDa and 100 kDa markers and blocked in TBST/5% milk. The middle portion (50–100 kDa) was incubated with anti-NUAK1 antibody (1:500 TBST/5% milk), and the bottom portion (<50 kDa) was incubated with anti-GAPDH antibody (1:5,000 TBST/5% milk) at 4 °C overnight. After washing, these blots were incubated with an anti-rabbit HRP secondary antibody (diluted 1:1,000 in TBST with 5% milk) at room temperature for 1 hour. The top portion (>100 kDa) was incubated with the pre-cleared anti-MYPT1 pSer445 antibody overnight at 4 °C, washed, and subsequently incubated with an anti-sheep HRP secondary antibody (diluted 1:1,000 in TBST/5% milk) at room temperature for 1 hour. After chemiluminescence detection, the MYPT1 pSer445 blot was gently washed for 1 hour in TBS, re-blocked, and re-probed for 1 hour with total anti-MYPT1 antibody (1:200 in TBST/5% milk). The blot was then incubated for 1 hour with an anti-mouse HRP secondary antibody (1:1,000 TBST/5% milk), with all incubation steps performed at room temperature. For signal quantification, ImageStudio (version 6.0.0.28) was used to measure band intensity for MYPT1 (pSer445), total MYPT1, and GAPDH. pSer445 intensity was normalized to total MYPT1 from the same sample. Signals for each blot were normalized to mock vehicle control.

### Generation of genetically modified primary human CD34^+^ HSPCs

Human bone-marrow-derived CD34^+^ hematopoietic stem cells (HSPCs) from de-identified donors (Stem Cell Technologies) were cultured as previously described ^72,28^. To generate NUAK1-null cells, two sgRNAs targeting exon 1 of human NUAK1 (Synthego; NUAK1_CR1: 5’-UGAUGCCGCUUCACCCCGUG-3’ and NUAK1_CR2: 5’-CACUCGGCCAGAAAACCUCU-3’) were complexed with recombinant Cas9-NLS protein (Berkeley Macrolab) to form ribonucleoprotein complexes (RNPs). On day 2 of culture, RNPs were delivered into HSPCs via nucleofection (Lonza, Program EO-100). On day 7, the cells were induced to differentiate down the erythroid lineage as previously described ^72,28^. To quantify knockout efficiency, genomic DNA was isolated from cells on days 8 and 15 of differentiation. The targeted locus was PCR-amplified using primers N1KO_F1 (5’-TCCCCTCCCTCCTGCAAACACC-3’) and N1KO_R1 (5’-CTGAGTCACGGGGTGCTGGAGA-3’). Indel frequency was quantified from the Sanger sequencing data of the resulting amplicons using the ICE Analysis tool (Synthego).

### Kinexus antibody microarrays

Kinexus 900P and 1325P phospho-antibody microarray kits were purchased from Kinexus (Vancouver, Canada). WT and CD44-null day 19 cRBCs were incubated with 2 µM recombinant EBA-175, generated as previously described ^73,74^, or PBS for 30 minutes in buffer HBSS. For the arrays performed with *P.falciparum-*stimulated cRBCs, cells were incubated with purified rupturing schizonts (10% parasitemia and 0.5% hematocrit) for 3 hours and harvested (3.7 × 10^6^ schizonts/3.7 × 10^7^ CD44-null cRBCs; 5 × 10^6^ schizonts/5 × 10^7^ WT cRBC). Cells were washed in ice-cold PBS and protein extracts were prepared as per manufacturer’s instructions with Kinexus Lysis Buffer supplemented with protease inhibitors, 1 mM DTT, and chemical cleavage reagents TCEP and NTCB. The microarray scanning, quantification, and analysis were performed by Kinexus. For Kinexus microarray heatmaps, the data were sorted by the unique antibody protein codes and only phospho-specific antibodies were considered. Tiles were arranged in a grid and colored by % Change from mock using R (version 4.4.2) and ggplot2 (version 3.5.1).

### Immunofluorescence assays

Magnet-purified schizonts were incubated in 25 nM ML-10 for 3.5 hours, washed extensively in warm media, and then incubated with donor RBCs at 2% hematocrit at 10-20% parasitemia in the presence of 1 µM CytD and either 10 µM HTH-01-015 or mock vehicle control (0.1% DMSO) in cRPMI. After 45 minutes, 120 µL of each sample was added to 1 mL of fix solution (4% PFA, 0.0075% Glutaraldehyde, PBS) for 20 minutes at room temperature and processed for IFA as previously described, with some modifications ^75^. Samples were washed, allowed to settle on glass cover slips, and then permeabilized with 0.1% Triton X-100 in PBS for 8 minutes. After washing, they were blocked with PBS/3% BSA for one hour at room temperature and then incubated overnight with 1:500 of anti-PfRAP1 209.3 ^70^ in PBS/3% BSA at 4_°_C. After washing, samples were incubated in 1:2,000 of Alexafluor 647 anti-mouse secondary antibody in PBS/3% BSA, incubated in 1:5,000 Hoechst (Invitrogen, #H3570), and mounted to a glass slide with Fluoromount-G (ThermoFisher, #00-4958-02). Images of attached merozoites were captured with a 100X objective on a Keyence BZ-X700 microscope. For quantification of rhoptry release, ∼80 images of attached merozoites in each condition obtained in two independent experiments were scored blindly for the presence or absence of discharged RAP1 signal within the RBC.

### Kinobead competition experiments and nLC-MS Analysis

For human kinome selectivity profiling of inhibitors, a 1:1 mixture of lysate of the hepatocyte line Friendship of China and US (FOCUS) and the T-lymphocyte line Jurkat was used. Briefly, cells were rinsed twice with ice-cold PBS and either pelleted (Jurkat, suspension) before being lysed in modified RIPA buffer (50 mM Tris-HCl, 150 mM NaCl, 1% Nonidet P40 Substitute (v/v) (Sigma), 0.25% sodium deoxycholate (w/v), 1 mM EDTA, 10 mM NaF, 5% glycerol (v/v), pH 7.8), containing HALT protease inhibitor cocktail (100x, ThermoFisher), or lysed in modified RIPA buffer on the dish and harvested using a cell scraper (FOCUS, adherent). Lysates were vortexed five times intermittently at 21,000 x g for 5s each while being kept on ice and lysates clarified by centrifugation (21,000 x g, 20 min, 4°C). Protein content was determined using the Pierce 660 nm assay (Thermo), protein concentrations adjusted to 4 mg/mL using modified RIPA buffer, and lysates mixed 1:1.

For producing *P. falciparum* lysate, trophozoite and schizont stage parasites were purified by MACS and incubated for five minutes in ice-cold 0.15% saponin in PBS with protease and phosphate inhibitors, to separate the parasites from the red blood cell fraction. The parasite fraction was washed twice in ice-cold PBS, and the pellet was lysed in modified RIPA buffer, and the lysates vortexed, clarified, and the protein content measured as described above. Lysate protein concentration was adjusted to 4 mg/mL using modified RIPA buffer. All operations were performed on wet ice. Kinome selectivity profiling of inhibitors was performed using an in-house developed Kinobead AP-MS competition assay ^46,76^.

Peptide samples from human kinome profiling were analyzed on a Thermo Orbitrap Fusion Lumos system. Peptide samples were separated on an EASY-nLC 1200 System (Thermo Fisher Scientific) using 20 cm long fused silica capillary columns (100 µm ID, laser pulled in-house with Sutter P-2000, Novato CA) packed with 3 μm 120 Å reversed phase C18 beads (Dr. Maisch, Ammerbuch, DE). The LC gradient was 90 min long with 5−35% B at 300 nL/min. LC solvent A was 0.1% (v/v) aq. acetic acid and LC solvent B was 20% 0.1% (v/v) acetic acid, 80% acetonitrile. MS data was collected with a Thermo Fisher Scientific Orbitrap Fusion Lumos. Data-dependent analysis was applied using Top15 selection with CID fragmentation. MS raw files were computed using MaxQuant/Andromeda v1.5.2.8 with the label-free quantification (LFQ) feature enabled and using the human FASTA (UP000005640) downloaded from Uniprot on July 22, 2015, as described previously ^77,78^.

Peptides from *P. falciparum* kinome profiling were analyzed on a Bruker nanoElute 2 – timsTOF Pro 2 nanoLC-MS system using the diaPASEF method with label-free quantification (LFQ) ^76,79^. MS raw files were searched using DIA-NN v2.0.1 in library-free mode. To distinguish *P. falciparum* from human kinases, both the *P. falciparum* (isolate 3D7) FASTA (UP000001450) downloaded from the Uniprot database on July 12, 2024 and the human FASTA (UP000005640) downloaded from Uniprot on July 14, 2023 were searched. DIA-NN output files processed using Perseus v2.0.10.0 ^80,81^.

### Split-luciferase kinase assays

Kinase selectivity profiling was performed by Luceome Biotechnologies (Tucson, AZ) using their split-luciferase KinaseSeeker™ competition binding assay ^56^. Initially, HTH-01-015 and WZ4003 were screened in duplicate at 10 µM against a custom panel of 11 *P. falciparum* kinases alongside human NUAK1 and NUAK2. For each assay, a Cfluc-tagged kinase and a Fos-Nfluc fusion protein were expressed in a cell-free rabbit reticulocyte lysate system. Lysate containing the split-luciferase components was incubated with the inhibitor or DMSO (vehicle control) for 2 hours in the presence of a kinase-specific probe. After incubation, a luciferin assay reagent was added, and luminescence was immediately measured as per manufacturer’s instructions. The percent inhibition was calculated using the formula: % Inhibition = [(Control Luminescence – Sample Luminescence) / Control Luminescence] × 100. The % Activity Remaining was reported as 100 - % Inhibition. Based on the initial screening results, IC₅₀ values were subsequently determined for select kinases by Luceome. For these assays, HTH-01-015 and WZ4003 were serially diluted and tested in duplicate across an eight-point curve.

### Generation of NUAK1-overexpressing BEL-A cells

Human NUAK1 cDNA was ordered from GenScript (#OHu17138) and cloned into pLVX-EF1alpha-SARS-CoV-2-E-2xStrep-IRES-Puro (Addgene plasmid #141385, a gift from Nevan Krogan) by Gibson cloning (New England Biolabs #E5510S) with forward primer, 5’-GTGTCGTGAGGATCTATTTCCGGTGCATGGAAGGGGCCGCCGCGCCTGTG-3’, and reverse primer, 5’-AGGGGGGGGGGAGGGAGAGGGGCGGATCACTTATCGTCGTCATCCTTGTAATCGTTG AGC-‘3. Lentivirus was produced in HEK293T cells (ATCC) using standard methods. Lentivirus transductions were performed in 24 well plates with 1×10^6^ BEL-A cells in 1.2 mL StemSpan, 800 µL of the lentivirus, and 8 µL of 1mg/ml polybrene. Spinoculation was performed by centrifugation at 1,000 xG for 2 hours at 30_°_C. Cells were then washed and resuspended in complete StemSpan at 1 × 10^5^ cell/mL at 37_°_C in 5% CO_2_. 1 µg/mL puromycin was used to select for transduced cells.

### *P. falciparum* invasion assays in BEL-A-derived orthochromatic erythroblasts

For *P. falciparum* invasion assays in BEL-A-derived erythroblasts, WT or NUAK1-overexpressing BEL-A cells were differentiated for 8 days, and orthochromatic erythroblasts were isolated using a Percoll gradient. These cells were incubated with purified, late-stage *P. falciparum* schizonts in duplicate in the presence of 1.25 µM HTH-01-015, 5 µM HTH-01-015, or a mock control (0.05% DMSO). All assays were conducted in a 96-well plate in a 100 µL volume containing cRPMI, 0.2% hematocrit, and a 10% initial parasitemia. Cytospin slides were prepared the next day (16-18 hours post-incubation), and the ring-stage parasitemia was determined by blinded counting of at least 2,000 cells per slide by two independent researchers, which were averaged to yield the reported parasitemia for each slide. Two independent biological replicates were performed for each drug concentration, with two technical replicates per condition.

### Data Analysis

Flow cytometry data was processed in FlowJo (version 10.10.0). Invasion assay plots, growth assay plots, IC_50_ calculations, and statistical analyses were performed using GraphPad Prism (version 10). Statistical tests included two-tailed unpaired Student’s t-test, one-way analysis of variance (ANOVA), and two-way ANOVA. IC₅₀ values were determined by fitting dose-response data to a four-parameter non-linear regression curve. Kinobead pulldown data was analyzed as described in the relevant methods section and plotted using R (version 4.4.2). For differential expression analysis of MS-based proteomics data we used two-tailed two-sample Student’s t-tests with Benjamini-Hochberg correction for multiple hypothesis testing (*N*=4). To search for a NUAK1 ortholog in *P. falciparum*, we used the PlasmoDB BLASTp search tool with the human NUAK1 amino acid sequence.

## ACKNOWLEDGEMENTS

The authors thank Steven Pelech from Kinexus, Reena Zutshi from Luceome, and Melanie Espiritu, Chhaminder Kaur, and Tianjian Lin from the Egan Lab for technical assistance. The authors are grateful to Tim Satchwell and Jan Frayne for the BEL-A cell line, James Hastie at the MRC for the MYPT1 antibodies, Anthony Holder for anti-RAP1/2 monoclonal antibody 209.3, Dave Richard for D10PfPHG, and Simon Osborne and LifeArc for providing ML-10 (compound MRT00207065). The unmodified BEL-A cell line was created by Professor Jan Frayne, Professor David Anstee and Dr. Kongtana Trakarnsanga with funding from the Wellcome Trust (grant numbers 087430/Z/08 and 102610), NHS Blood and Transplant and Department of Health (England). We thank the Stanford Blood Center for the supply of human blood for parasite culture, and PlasmoDB hosted by the VEuPathDB resource center for providing *Plasmodium* genomic resources. This work used an EASY-nLC1200 UHPLC and Thermo Scientific Orbitrap Fusion Lumos Tribrid mass spectrometer purchased with funding from a National Institutes of Health SIG grant S10OD021502. The authors are grateful to Debopam Chakrabarti, John Boothroyd, Shirit Einav, Marwah Karim, Nathanael Gray, Priscilla Yang, David Schneider, Sourav Banerjee, and members of Egan, Boothroyd, and Mattias Garten Labs for helpful discussions. This work was supported by funding from the NIH National Heart Lung and Blood Institute Grants R01HL166249 and DP2HL13718601 (ESE), as well as a Bridge Grant award from the American Society of Hematology (ESE). DJN was funded by a Stanford Graduate Smith Fellowship. ESE is a Chan Zuckerberg Biohub-San Francisco investigator and a Tashia and John Morgridge Endowed Faculty Scholar of the Stanford Maternal and Child Health Research Institute. This research was supported [in part] by the Intramural Research Program of the National Institutes of Health (NIH). The contributions of the NIH author(s) are considered Works of the United States Government. The findings and conclusions presented in this paper are those of the author(s) and do not necessarily reflect the views of the NIH or the U.S. Department of Health and Human Services.

## Author contributions

D.J.N., C.Y.K., M.G. and E.S.E. conceived of the study; D.J.N., C.Y.K., K.M., M.G.R., B.B., N.D.S., M.G.G. and E.S.E performed experiments and analyzed data; N.H.T., C.D., S-E.O., M.G.G. and E.S.E. provided supervision and guidance; D.J.N, and E.S.E. wrote the original manuscript draft; N.H.T., C.D., S-E.O., M.G.G., and E.S.E. reviewed and edited the manuscript. All authors approved the final manuscript.

## Extended data

Extended Data Figures 1-4

Extended Data Table 1: Proteomics Data: Human and *Plasmodium* Kinome Profiling

Extended Data Videos 1-4

## REFERENCES

1. World Health Organization. World Malaria Report 2024: Addressing Inequity in the Global Malaria Response. (2024).

2. Haldar, K., Bhattacharjee, S. & Safeukui, I. Drug resistance in Plasmodium. Nat Rev Microbiol 16, 156–170 (2018).

3. Ariey, F. et al. A molecular marker of artemisinin-resistant Plasmodium falciparum malaria. Nature 505, 50–55 (2014).

4. Mita, T. et al. Limited Geographical Origin and Global Spread of Sulfadoxine-Resistant dhps Alleles in Plasmodium falciparum Populations. The Journal of Infectious Diseases 204, 1980–1988 (2011).

5. Ashley, E. A. et al. Spread of Artemisinin Resistance in *Plasmodium falciparum* Malaria. N Engl J Med 371, 411–423 (2014).

6. Balikagala, B. et al. Evidence of Artemisinin-Resistant Malaria in Africa. The New England Journal of Medicine 385, 1163–1171 (2021).

7. Thriemer, K. et al. Delayed parasite clearance after treatment with dihydroartemisinin-piperaquine in Plasmodium falciparum malaria patients in central Vietnam. Antimicrobial Agents and Chemotherapy 58, 7049–7055 (2014).

8. Birnbaum, J. et al. A Kelch13-defined endocytosis pathway mediates artemisinin resistance in malaria parasites. Science 367, 51–59 (2020).

9. Okombo, J. & Fidock, D. A. Towards next-generation treatment options to combat Plasmodium falciparum malaria. Nat Rev Microbiol 23, 178–191 (2025).

10. Zumla, A. et al. Host-directed therapies for infectious diseases: current status, recent progress, and future prospects. The Lancet Infectious Diseases 16, e47–63 (2016).

11. Glennon, E. K. K., Dankwa, S., Smith, J. D. & Kaushansky, A. Opportunities for Host-targeted Therapies for Malaria. Trends in Parasitology 34, 843–860 (2018).

12. Duffey, M. et al. Combating antimicrobial resistance in malaria, HIV and tuberculosis. Nat Rev Drug Discov 23, 461–479 (2024).

13. Wei, L. et al. Host-directed therapy, an untapped opportunity for antimalarial intervention. Cell reports. Medicine 2, 100423 (2021).

14. Jeong, E.-K., Lee, H.-J. & Jung, Y.-J. Host-Directed Therapies for Tuberculosis. Pathogens 11, 1291 (2022).

15. Kumar, N. et al. Host-Directed Antiviral Therapy. Clinical Microbiology Reviews 33, 10.1128/cmr.00168-19 (2020).

16. Vijayan, K. et al. Host-targeted Interventions as an Exciting Opportunity to Combat Malaria. Chemical Reviews 121, 10452–10468 (2021).

17. Lelliott, P. M., McMorran, B. J., Foote, S. J. & Burgio, G. The influence of host genetics on erythrocytes and malaria infection: is there therapeutic potential? Malaria Journal 14, 289 (2015).

18. Lopaticki, S. et al. Reticulocyte and Erythrocyte Binding-Like Proteins Function Cooperatively in Invasion of Human Erythrocytes by Malaria Parasites. Infect Immun 79, 1107–1117 (2011).

19. Weiss, G. E. et al. Revealing the Sequence and Resulting Cellular Morphology of Receptor-Ligand Interactions during Plasmodium falciparum Invasion of Erythrocytes. PLOS Pathogens 11, e1004670 (2015).

20. Cowman, A. F., Tonkin, C. J., Tham, W.-H. & Duraisingh, M. T. The Molecular Basis of Erythrocyte Invasion by Malaria Parasites. Cell Host & Microbe 22, 232–245 (2017).

21. Tolia, N. H., Enemark, E. J., Sim, B. K. L. & Joshua-Tor, L. Structural basis for the EBA-175 erythrocyte invasion pathway of the malaria parasite Plasmodium falciparum. Cell 122, 183– 193 (2005).

22. Salinas, N. D., Paing, M. M. & Tolia, N. H. Critical Glycosylated Residues in Exon Three of Erythrocyte Glycophorin A Engage Plasmodium falciparum EBA-175 and Define Receptor Specificity. mBio 5, 10.1128/mbio.01606-14 (2014).

23. Koch, M. et al. Plasmodium falciparum erythrocyte-binding antigen 175 triggers a biophysical change in the red blood cell that facilitates invasion. Proceedings of the National Academy of Sciences of the United States of America 114, 4225–4230 (2017).

24. Sisquella, X. et al. Plasmodium falciparum ligand binding to erythrocytes induce alterations in deformability essential for invasion. eLife 6, (2017).

25. Zuccala, E. S. et al. Quantitative phospho-proteomics reveals the Plasmodium merozoite triggers pre-invasion host kinase modification of the red cell cytoskeleton. Sci Rep 6, 19766 (2016).

26. Paing, M. M. et al. Shed EBA-175 mediates red blood cell clustering that enhances malaria parasite growth and enables immune evasion. Elife 7, e43224 (2018).

27. Egan, E. S. et al. A forward genetic screen identifies erythrocyte CD55 as essential for Plasmodium falciparum invasion. Science 348, 711–714 (2015).

28. Baro, B., et al. *Plasmodium falciparum* exploits CD44 as a coreceptor for erythrocyte invasion. Blood 142, 2016–2028 (2023).

29. Adderley, J. D. et al. Analysis of erythrocyte signalling pathways during Plasmodium falciparum infection identifies targets for host-directed antimalarial intervention. Nature Communications 11, 4015 (2020).

30. Suzuki, A. et al. Identification of a Novel Protein Kinase Mediating Akt Survival Signaling to the ATM Protein *. Journal of Biological Chemistry 278, 48–53 (2003).

31. Kusakai, G., Suzuki, A., Ogura, T., Kaminishi, M. & Esumi, H. Strong association of ARK5 with tumor invasion and metastasis. J Exp Clin Cancer Res 23, 263–268 (2004).

32. Cui, J. et al. Overexpression of ARK5 is associated with poor prognosis in hepatocellular carcinoma. Tumour Biol 34, 1913–1918 (2013).

33. Port, J. et al. Colorectal Tumors Require NUAK1 for Protection from Oxidative Stress. Cancer discovery 8, 632–647 (2018).

34. Skalka, G. L., Whyte, D., Lubawska, D. & Murphy, D. J. NUAK: never underestimate a kinase. Essays in biochemistry 10.1042/EBC20240005 2024) doi:10.1042/EBC20240005.

35. Zagórska, A. et al. New roles for the LKB1-NUAK pathway in controlling myosin phosphatase complexes and cell adhesion. Science Signaling 3, ra25 (2010).

36. Golkowski, M. et al. Pharmacoproteomics Identifies Kinase Pathways that Drive the Epithelial-Mesenchymal Transition and Drug Resistance in Hepatocellular Carcinoma. Cell Syst 11, 196–207.e7 (2020).

37. Lizcano, J. M. et al. LKB1 is a master kinase that activates 13 kinases of the AMPK subfamily, including MARK/PAR-1. The EMBO Journal 23, 833–843 (2004).

38. Banerjee, S. et al. Characterization of WZ4003 and HTH-01-015 as selective inhibitors of the LKB1-tumour-suppressor-activated NUAK kinases. The Biochemical Journal 457, 215– 225 (2014).

39. Trakarnsanga, K. et al. An immortalized adult human erythroid line facilitates sustainable and scalable generation of functional red cells. Nat Commun 8, 1–7 (2017).

40. Baker, D. A. et al. A potent series targeting the malarial cGMP-dependent protein kinase clears infection and blocks transmission. Nat Commun 8, 430 (2017).

41. Paul, A. S., Egan, E. S. & Duraisingh, M. T. Host-parasite interactions that guide red blood cell invasion by malaria parasites. Current Opinion in Hematology 22, 220–226 (2015).

42. Volz, J. C. et al. Essential Role of the PfRh5/PfRipr/CyRPA Complex during Plasmodium falciparum Invasion of Erythrocytes. Cell Host & Microbe 20, 60–71 (2016).

43. Weiss, G. E., Crabb, B. S. & Gilson, P. R. Overlaying Molecular and Temporal Aspects of Malaria Parasite Invasion. Trends in Parasitology 32, 284–295 (2016).

44. Scally, S. W. et al. PCRCR complex is essential for invasion of human erythrocytes by Plasmodium falciparum. Nat Microbiol 7, 2039–2053 (2022).

45. Bantscheff, M. et al. Quantitative chemical proteomics reveals mechanisms of action of clinical ABL kinase inhibitors. Nat Biotechnol 25, 1035–1044 (2007).

46. Golkowski, M. et al. Kinobead and Single-Shot LC-MS Profiling Identifies Selective PKD Inhibitors. Journal of Proteome Research 16, 1216–1227 (2017).

47. Uitdehaag, J. C. et al. A guide to picking the most selective kinase inhibitor tool compounds for pharmacological validation of drug targets. British Journal of Pharmacology 166, 858– 876 (2012).

48. van Bergen, W., Nederstigt, A. E., Heck, A. J. R. & Baggelaar, M. P. Site-Specific Competitive Kinase Inhibitor Target Profiling Using Phosphonate Affinity Tags. Molecular & Cellular Proteomics 24, 100906 (2025).

49. Kovackova, S. et al. Selective Inhibitors of Cyclin G Associated Kinase (GAK) as Anti-Hepatitis C Agents. Journal of Medicinal Chemistry 58, 3393–3410 (2015).

50. Ward, P., Equinet, L., Packer, J. & Doerig, C. Protein kinases of the human malaria parasite Plasmodium falciparum: the kinome of a divergent eukaryote. BMC Genomics 5, 1–19 (2004).

51. Adderley, J. & Doerig, C. Comparative analysis of the kinomes of Plasmodium falciparum, Plasmodium vivax and their host Homo sapiens. BMC Genomics 23, 237 (2022).

52. Adderley, J., Williamson, T. & Doerig, C. Parasite and Host Erythrocyte Kinomics of Plasmodium Infection. Trends in Parasitology 37, 508–524 (2021).

53. Solyakov, L. et al. Global kinomic and phospho-proteomic analyses of the human malaria parasite Plasmodium falciparum. Nat Commun 2, 1–12 (2011).

54. Miranda-Saavedra, D., Gabaldón, T., Barton, G. J., Langsley, G. & Doerig, C. The kinomes of apicomplexan parasites. Microbes and Infection 14, 796–810 (2012).

55. Vidadala, R. S. R. et al. 7 H-Pyrrolo[2,3-d]pyrimidin-4-amine-Based Inhibitors of Calcium-Dependent Protein Kinase 1 Have Distinct Inhibitory and Oral Pharmacokinetic Characteristics Compared with 1 H-Pyrazolo[3,4-d]pyrimidin-4-amine-Based Inhibitors. ACS Infect Dis 4, 516–522 (2018).

56. Bohmer, M. J. et al. Human Polo-like Kinase Inhibitors as Antiplasmodials. ACS Infectious Diseases 9, 1004–1021 (2023).

57. Jester, B. W. et al. A Coiled-Coil Enabled Split-Luciferase Three-Hybrid System: Applied Toward Profiling Inhibitors of Protein Kinases. J Am Chem Soc 132, 11727–11735 (2010).

58. Ganter, M. et al. Plasmodium falciparum CRK4 directs continuous rounds of DNA replication during schizogony. Nat Microbiol 2, 1–9 (2017).

59. Zhang, M. et al. Uncovering the essential genes of the human malaria parasite Plasmodium falciparum by saturation mutagenesis. Science 360, eaap7847 (2018).

60. Satchwell, T. J. et al. Genetic manipulation of cell line derived reticulocytes enables dissection of host malaria invasion requirements. Nat Commun 10, 3806 (2019).

61. Angulo-Urarte, A. et al. Endothelial cell rearrangements during vascular patterning require PI3-kinase-mediated inhibition of actomyosin contractility. Nat Commun 9, 1–16 (2018).

62. Schor, S. & Einav, S. Combating Intracellular Pathogens with Repurposed Host-Targeted Drugs. ACS Infect. Dis. 4, 88–92 (2018).

63. Thom, R. E. & D’Elia, R. V. Future applications of host direct therapies for infectious disease treatment. Front. Immunol. 15, (2024).

64. Adderley, J. & Grau, G. E. Host-directed therapies for malaria: possible applications and lessons from other indications. Current Opinion in Microbiology 71, 102228 (2022).

65. Boulet, C. et al. Red Blood Cell BCL-xL Is Required for Plasmodium falciparum Survival: Insights into Host-Directed Malaria Therapies. Microorganisms 10, (2022).

66. Kesely, K. R., Pantaleo, A., Turrini, F. M., Olupot-Olupot, P. & Low, P. S. Inhibition of an Erythrocyte Tyrosine Kinase with Imatinib Prevents Plasmodium falciparum Egress and Terminates Parasitemia. PLOS ONE 11, e0164895 (2016).

67. Pantaleo, A. et al. Syk inhibitors interfere with erythrocyte membrane modification during P falciparum growth and suppress parasite egress. Blood 130, 1031–1040 (2017).

68. Chien, H. D. et al. Imatinib augments standard malaria combination therapy without added toxicity. The Journal of Experimental Medicine 218, (2021).

69. Banerjee, S. et al. Interplay between Polo kinase, LKB1-activated NUAK1 kinase, PP1\betaMYPT1 phosphatase complex and the SCF\betaTrCP E3 ubiquitin ligase. The Biochemical Journal 461, 233–245 (2014).

70. Wilson, D. W., Crabb, B. S. & Beeson, J. G. Development of fluorescent Plasmodium falciparum for in vitro growth inhibition assays. Malaria Journal 9, 152 (2010).

71. Howell, S. A. et al. Distinct mechanisms govern proteolytic shedding of a key invasion protein in apicomplexan pathogens. Molecular Microbiology 57, 1342–1356 (2005).

72. Kaneko-Kawano, T. et al. Dynamic Regulation of Myosin Light Chain Phosphorylation by Rho-kinase. PLOS ONE 7, e39269 (2012).

73. Shakya, B., Patel, S. D., Tani, Y. & Egan, E. S. Erythrocyte CD55 mediates the internalization of Plasmodium falciparum parasites. eLife 10, (2021).

74. Chen, E., Paing, M. M., Salinas, N., Sim, B. K. L. & Tolia, N. H. Structural and Functional Basis for Inhibition of Erythrocyte Invasion by Antibodies that Target Plasmodium falciparum EBA-175. PLOS Pathogens 9, e1003390 (2013).

75. Salinas, N. D. & Tolia, N. H. A quantitative assay for binding and inhibition of *Plasmodium falciparum* Erythrocyte Binding Antigen 175 reveals high affinity binding depends on both DBL domains. Protein Expression and Purification 95, 188–194 (2014).

76. Tonkin, C. J. et al. Localization of organellar proteins in *Plasmodium falciparum* using a novel set of transfection vectors and a new immunofluorescence fixation method. Molecular and Biochemical Parasitology 137, 13–21 (2004).

77. Woods, K. et al. diaPASEF-Powered Chemoproteomics Enables Deep Kinome Interaction Profiling. J Proteome Res 24, 4463–4477 (2025).

78. Cox, J. et al. Accurate proteome-wide label-free quantification by delayed normalization and maximal peptide ratio extraction, termed MaxLFQ. Mol Cell Proteomics 13, 2513–2526 (2014).

79. Golkowski, M. et al. Multiplexed kinase interactome profiling quantifies cellular network activity and plasticity. Mol Cell 83, 803–818.e8 (2023).

80. Meier, F. et al. diaPASEF: parallel accumulation-serial fragmentation combined with data-independent acquisition. Nat Methods 17, 1229–1236 (2020).

81. Demichev, V., Messner, C. B., Vernardis, S. I., Lilley, K. S. & Ralser, M. DIA-NN: neural networks and interference correction enable deep proteome coverage in high throughput. Nat Methods 17, 41–44 (2020).

82. Tyanova, S. et al. The Perseus computational platform for comprehensive analysis of (prote)omics data. Nat Methods 13, 731–740 (2016).

