## Supplemental Figures for "*Plasmodium falciparum* exploits NUAK1 to establish infection in human erythrocytes"

**a**

WT cRBCs: EBA-175-stimulated vs mock

CD44-null cRBCs: EBA175-stimulated vs mock

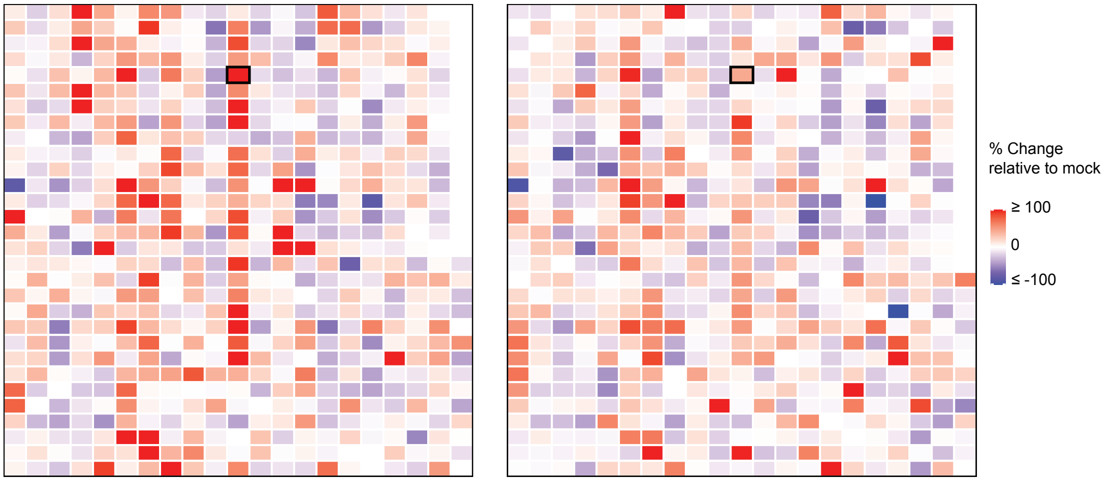


**b**

WT cRBCs: *Pf*3D7-stimulated vs mock

CD44-null cRBCs: *Pf*3D7-stimulated vs mock

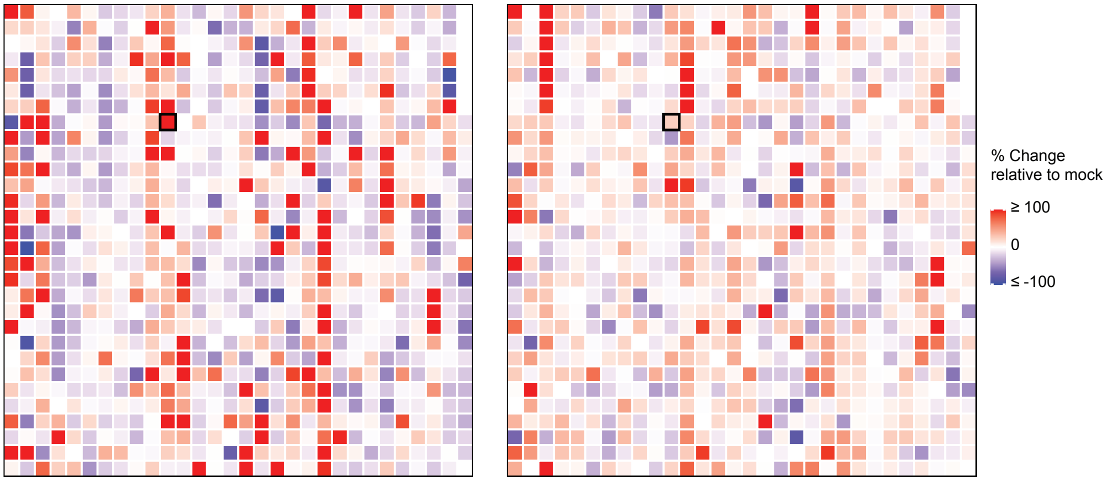


c

| Target Protein Name | Phosphosite | % Change relative to mock | |
|---|---|---|---|
| | | EBA-175-stimulated WT | EBA-175-stimulated CD44-null |
| Adducin a/g | S726 | 73 | 32 |
| Bmx | Y40 | 77 | 27 |
| CDK12 | T893 | 75 | 91 |
| CDK2 | T160 | 640 | 706 |
| CHK2 | T68 | 108 | 117 |
| eIF4G | S1106 | -75 | -33 |
| FLT3 | Y842 | 54 | 39 |
| FYN | Y531 | 60 | 11 |
| MDM2 | S166 | -52 | -43 |
| MKK7 | T275 | 81 | 28 |
| MOK | T159+Y161 | 62 | 45 |
| NDR1 | S281+T282 | 72 | 47 |
| NEK2 | T170+S171 | 109 | 93 |
| NUAK1 | T211 | 107 | 43 |
| OSR1 | T185 | 51 | 14 |
| Raf1 (c-Raf) | S296 | 5745 | -6 |
| Raf1 (c-Raf) | S259 | 100052 | -30 |
| SIT | Y90 | 51 | 34 |
| SMC1 | S957 | 247 | 158 |
| SRC | Y419 | -56 | -44 |
| TAK1 | S439 | 104 | 46 |

d

| Target Protein Name | Phosphosite | % Change relative to mock | |
|---|---|---|---|
| | | *Pf*3D7-stimulated WT | *Pf*3D7-stimulated CD44-null |
| Adducin a/g | S726 | 54 | 53 |
| n.d. | | | |
| CDK12 | T893 | -5 | -18 |
| CDK2 | T160 | 26 | -38 |
| CHK2 | T68 | -11 | 2 |
| eIF4G | S1106 | 34 | -26 |
| FLT3 | Y842 | -19 | -5 |
| FYN | Y531 | -14 | 13 |
| MDM2 | S166 | 26 | -63 |
| MKK7 | T275 | 123 | -11 |
| MOK | T159+Y161 | 12 | -24 |
| NDR1 | S281+T282 | 6 | 26 |
| NEK2 | T170+S171 | 9 | 136 |
| NUAK1 | T211 | 109 | 24 |
| OSR1 | T185 | 105 | 19 |
| Raf1 (c-Raf) | S296 | 0 | 33 |
| Raf1 (c-Raf) | S259 | -21 | 35 |
| SIT | Y90 | -3 | 25 |
| n.d. | | | |
| SRC | Y419 | 248 | 13 |
| TAK1 | S439 | -15 | 33 |

**Extended Data Figure 1. Phospho-antibody microarray analysis of stimulated cRBCs.**

**a, b,** Heatmaps of phospho-antibody microarray data from wild-type (WT) and CD44-null cRBCs stimulated with either recombinant EBA-175 (**a**) or live *P. falciparum* 3D7 parasites (**b**). Data are displayed as the percent change from unstimulated controls (%CFC). The tile corresponding to the NUAK1 pThr211 antibody is outlined in black. **c, d,** Tables listing the top phosphorylated signals from the stimulation experiments. Lead signals were selected from the EBA-175 stimulation experiment (**c**) if they met the following criteria: %CFC ≥ 50; SUM of %Error Ranges < 0.85 × %CFC; and at least one Globally Normalized intensity value ≥ 1000. The table in **d** shows the corresponding data for this same set of proteins from the *P. falciparum* stimulation experiment. The NUAK1 pThr211 signal is highlighted in both tables. n.d., not done.

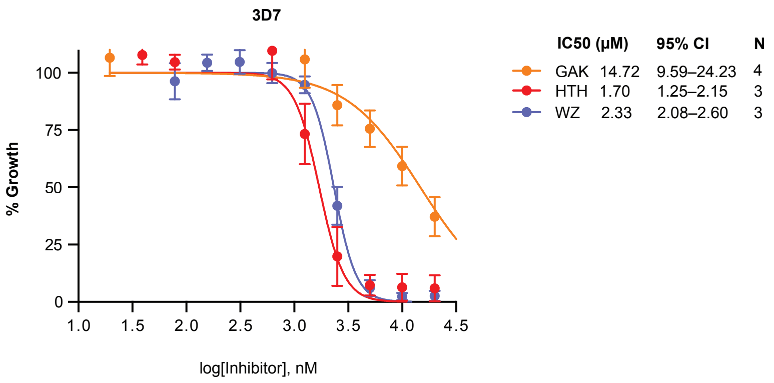


**Extended Data Figure 2. GAK inhibitor 12G has low antimalarial activity.**

Dose-response curves for *P. falciparum* strain 3D7 cultured for 72 hours with HTH-01-015, WZ4003, or GAK-specific inhibitor, 12G. Parasitemia was measured by flow cytometry and plotted relative to vehicle control. Data points are mean ± SEM. N = 3–4, as indicated.

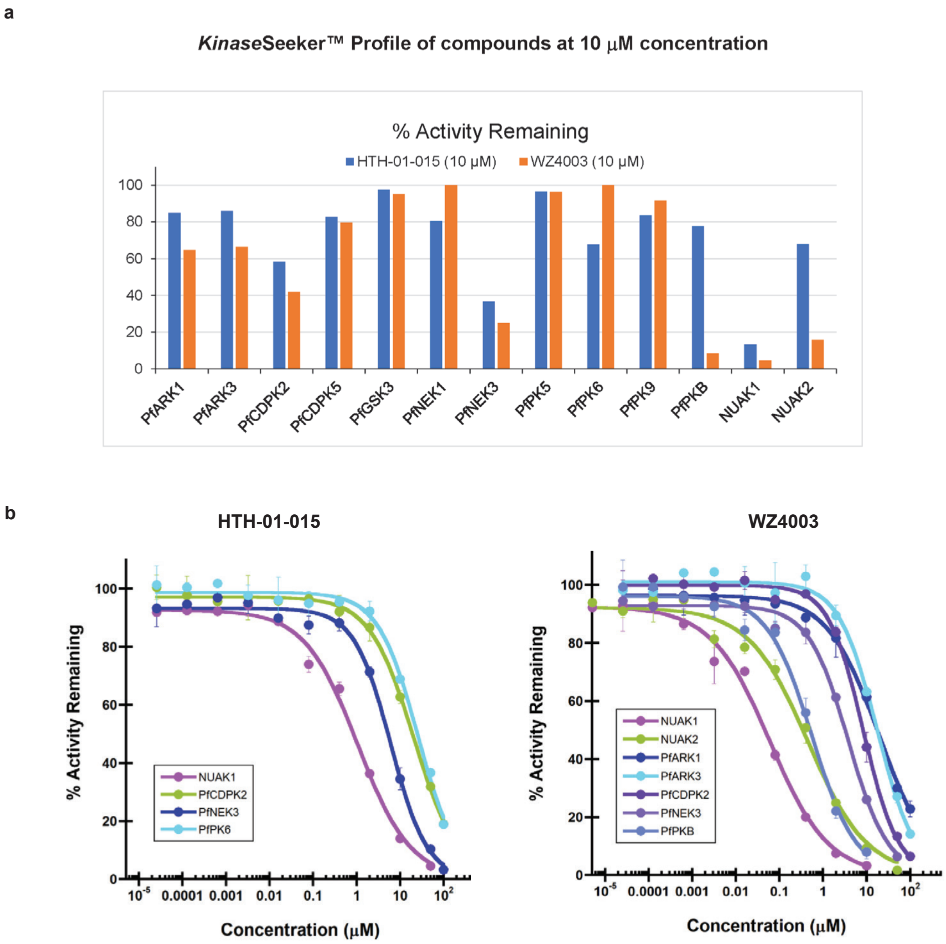


**Extended Data Figure 3. KinaseSeeker assay confirms high selectivity of HTH-01-015 and WZ4003 for human NUAK1, relative to *P. falciparum* kinases.**

**a,** Activity of luciferase-assembled kinases after 10 μM treatment of HTH-01-015 (blue) or WZ4003 (orange) for 11 Pf kinases and 2 human NUAK kinases. **b**, IC50 data for selected kinases for HTH-01-015 and WZ4003.

a

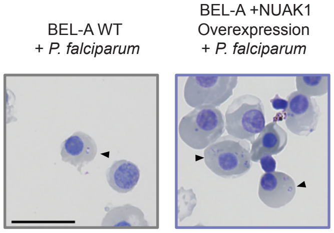


b

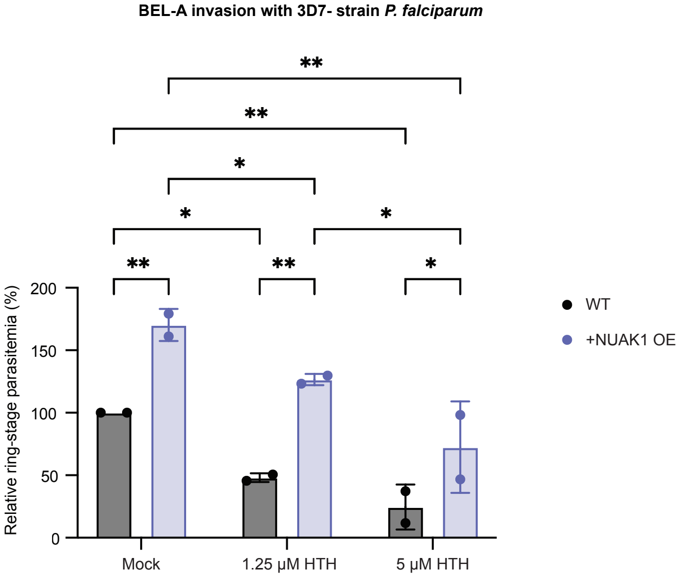


**Extended Data Figure 4. Increased *P. falciparum* invasion in NUAK1-overexpressing cells.**
**a,** Representative Giemsa-stained cytospin slides of WT BEL-A orthos and NUAK1-overexpressing BEL-A orthos infected with *P. falciparum* strain 3D7. Arrows indicate ring-stage parasites in infected cells. Scale bar = 20 μm. **b**, Parasitemias were measured by counting Giemsa-stained cytospin slides under a microscope, blinded. This is the same data as shown in Figure 5g, with parasitemias normalized to that of mock-treated WT orthos. Mean ± SD, N = 2. Statistical Analysis: Two-way ANOVA; ** $p < 0.01$, * $p < 0.05$.
